# Is structure based drug design ready for selectivity optimization?

**DOI:** 10.1101/2020.07.02.185132

**Authors:** Steven K. Albanese, John D. Chodera, Andrea Volkamer, Simon Keng, Robert Abel, Lingle Wang

## Abstract

Alchemical free energy calculations are now widely used to drive or maintain potency in small molecule lead optimization with a roughly 1 kcal/mol accuracy. Despite this, the potential to use free energy calculations to drive optimization of compound *selectivity* among two similar targets has been relatively unexplored in published studies. In the most optimistic scenario, the similarity of binding sites might lead to a fortuitous cancellation of errors and allow selectivity to be predicted more accurately than affinity. Here, we assess the accuracy with which selectivity can be predicted in the context of small molecule kinase inhibitors, considering the very similar binding sites of human kinases CDK2 and CDK9, as well as another series of ligands attempting to achieve selectivity between the more distantly related kinases CDK2 and ERK2. Using a Bayesian analysis approach, we separate systematic from statistical error and quantify the correlation in systematic errors between selectivity targets. We find that, in the CDK2/CDK9 case, a high correlation in systematic errors suggests free energy calculations can have significant impact in aiding chemists in achieving selectivity, while in more distantly related kinases (CDK2/ERK2), the correlation in systematic error suggests fortuitous cancellation may even occur between systems that are not as closely related. In both cases, the correlation in systematic error suggests that longer simulations are beneficial to properly balance statistical error with systematic error to take full advantage of the increase in apparent free energy calculation accuracy in selectivity prediction.

---

Free energy methods have proven useful in aiding structure-based drug design by driving the optimization or maintenance of potency in lead optimization. Alchemical free energy calculations allow for the prediction of ligand binding free energies, including all enthalpic and entropic contributions [1]. Advances in atomistic molecular mechanics simulations and free energy methodologies [2–5] have allowed free energy methods to reach a level of accuracy sufficient for predicting ligand potencies [6]. These methods have been applied prospectively to develop inhibitors for Tyk2 [7], Syk [8], BACE1 [9], GPCRs [10], and HIV protease [11]. A recent large-scale review of the use of FEP+ [12] to predict potency for 92 different projects and 3 021 compounds determined that predicted binding free energies had a median root mean squared error (RMSE) of 1.0 kcal/mol [13].

## Selectivity is an important consideration in drug design

In addition to potency, selectivity is an important property to consider in drug development, either in the pursuit of an inhibitor that is maximally selective [14, 15] or possesses a desired polypharmacology [16– 20]. Controlling selectivity can be useful not only in avoiding off-target toxicity (arising from inhibition of unintended targets) [21, 22], but also in avoiding on-target toxicity (arising from inhibition of the intended target) by selectively targeting disease mutations [23]. In either paradigm, considering the selectivity of a compound is complicated by the biology of the target. For example, kinases exist as nodes in complex signaling networks [24, 25] with feedback inhibition and cross-talk between pathways. Careful consideration of which off-targets are being inhibited can avoid off-target toxicity due to alleviating feedback inhibition and inadvertently reactivating the targeted pathway [24, 25] or the upregulation of a secondary pathway by alleviation of cross-talk inhibition [26, 27]. Off-target toxicity can also be caused by inhibiting unrelated targets, such as gefitinib, an EGFR inhibitor, inhibiting CYP2D6 [21] and causing hepatotoxicity in lung cancer patients. In a cancer setting, on-target toxicity can be avoided by considering the selectivity for the oncogenic mutant form of the kinase over the wild type form of the kinase [28–30], exemplified by a number of first generation EGFR inhibitors. Selective binding to multiple kinases can also lead to beneficial effects: Imatinib, initially developed to target BCR-Abl fusion proteins, is also approved for treating gastrointestinal stromal tumors (GIST) [31] due to its activity against receptor tyrosine kinase KIT.

## The use of physical modeling to predict selectivity is relatively unexplored

While engineering compound selectivity is important for drug discovery, the utility of free energy methods for predicting this selectivity with the aim of reducing the number of compounds that must be synthesized to achieve a desired selectivity profile has been relatively unexplored in published studies. If there is fortuitous cancellation of systematic errors for closely related systems, free energy methods may be much more accurate than expected given the errors made in predicting the potency for each individual target. Such systematic errors might arise from force field parameters uncertainty, force field parameters assignment, protein or ligand protonation state assignment, or even from systematic errors arising in the target experimental data, where for example poor solubility of a particular compound might lead to a spuriously poor reported binding affinity for that compound, among other effects.

Molecular dynamics and free energy calculations have been used extensively to investigate the biophysical origins of the selectivity of imatinib for Abl kinase over Src [32, 33] and within a family of non-receptor tyrosine kinases [34]. This work focused on understanding the role reorganization energy plays in the exquisite selectivity of imatinib for Abl over the highly related Src despite high similarity between the cocrystallized binding mode and kinase conformations, and touches on neither the evaluation of the accuracy of these methods nor their application to drug discovery on congeneric series of ligands. Previous work predicting the selectivity of three bromodomain inhibitors across the bromodomain family achieved promising accuracy for single target potency of roughly 1 kcal/mol, but does not explicitly evaluate any selectivity metrics [35] or quantify the correlation in the errors made in predicting affinities for each bromodomain. Previous work using FEP+ to predict selectivity between pairs of phosphodiesterases (PDEs) showed promising performance but did not evaluate correlation in the error made in predicting affinities for each PDE [36]

## Kinases are an important and particularly challenging model system for selectivity predictions

Kinases are a useful model system to work with for assessing the utility of free energy calculations to predict inhibitor selectivity in a drug discovery context. With the approval of imatinib for the treatment of chronic myelogenous leukemia in 2001, targeted small molecule kinase inhibitors (SMKIs) have become a major class of therapeutics in treating cancer and other diseases. Currently, there are 52 FDA-approved SMKIs [37], and it is estimated that kinase targeted therapies account for as much as 50% of current drug development [38], with many more compounds currently in clinical trials. While there have been a number of successful drug approvals, the current stable of FDA-approved kinase inhibitors targets only a small fraction of kinases implicated in disease, and the design of new selective kinase inhibitors for novel targets remains a significant challenge.

Achieving selective inhibition of kinases is quite challenging, as there are more than 518 protein kinases [39, 40] sharing a highly conserved ATP binding site that is targeted by the majority of SMKIs [41]. While kinase inhibitors have been designed to target kinase-specific sub-pockets and binding modes to achieve selectivity [42–47], previous work has shown that both Type I (binding to the active, DFG-in conformation) and Type II (binding to the inactive, DFG-out conformation) inhibitors are capable of achieving a range of selectivities [48, 49], often exhibiting significant binding to a number of other targets in addition to their primary target. Even FDA-approved inhibitors—often the result of extensive drug development programs—bind to a large number of off-target kinases [50]. Kinases are also targets of interest for developing polypharmacological compounds, or inhibitors that are specifically designed to inhibit multiple kinase targets. Resistance to MEK inhibitors in KRAS-mutant lung and colon cancer has been shown to be driven by ErbB3 upregulation [51], providing a rationale for dual MEK/ERBB family inhibitors. Similarly, combined MEK and VEGFR1 inhibition has been proposed as a combinatorial approach to treat KRAS-mutant lung cancer [52]. Developing inhibitors with a desired polypharmacology means navigating more complex selectivity profiles, presenting a problem where physical modeling has the potential to dramatically speedup drug discovery.

## The correlation coefficient measures how useful predictions are in achieving selectivity

Since the prediction of selectivity depends on predicting the change of affinities to two or more targets (or the change of affinities between pairs of related molecules for multiple targets), a spectrum of possibilities exists for how accurately selectivity can be predicted even when the error in predicting individual target affinities is fixed. In well-behaved kinase systems, for example, free energy calculation potency predictions have achieved root-mean-square of less than 1.0 kcal/mol [7, 12]. This residual error likely arises from a variety of contributions. Systematic contributions to the residual error may include forcefield parameterization deficiencies, protein and ligand protonation assignment errors, and discrepancies between the crystallographic construct protein and the assay construct protein. Likewise, unbiased contributions to the observed residual error likely arises from incompletely converged sampling. Lastly, it should not be forgotten that the target experimental value will have its own systematic and random errors.

In the best-case scenario, correlation in the systematic errors for predicting the interactions of a given ligand with two related protein targets might exactly cancel out, allowing selectivity to be predicted much more accurately than potency. On the other hand, if the uncorrelated random error dominates the residual error between two protein targets, predictions of selectivity will be *less accurate* than potency predictions. Real-world systems are likely to fall somewhere between these two extremes, and quantifying the *degree* to which error in multiple protein targets is correlated, its implications for the use of free energy calculations for prioritizing synthesis in the pursuit of selectivity, the ramifications for optimal calculation protocols, and rough guidelines governing which systems we might expect good selectivity prediction is the primary focus of this work.

In particular, in this work, we investigate the magnitude of the correlation (*ρ*) in error for the predicted binding free energy differences between related compounds (ΔΔ*G*_*ij*_) for two different targets, assessing the utility of alchemical free energy calculations for the prediction of selectivity. We employ state of the art relative free energy calculations [12, 13] to predict the selectivities of two different congeneric ligand series [53, 54], and construct simple numerical models that allow us to quantify the potential utility in selectivity optimization expected for different combinations of per target systematic errors and correlation coefficients. To make a realistic assessment of our confidence in this correlation coefficient derived from a limited number of experimental measurements, we develop a new Bayesian approach to quantify the uncertainty in the correlation coefficient in the predicted change in selectivity on ligand modification, incorporating all sources of uncertainty and correlation in the computation to separate statistical from systematic error. We find that in the closely related systems of CDK2 and CDK9, a high correlation of systematic errors suggests that free energy methods can have a significant impact on speeding up selectivity optimization. Even in the more distantly related case (CDK2/ERK2), correlation in the systematic errors allows free energy calculations to speedup selectivity optimization, suggesting that these methodologies can impact drug discovery even when comparing systems that are less closely related. We also present a model of the impact of per target statistical error at different levels of systematic error correlation, suggesting that it is worthwhile to expend more effort sampling in systems with high correlation.

## Results

### Alchemical free energy methods can be used to predict compound selectivity

While the potency of a ligand *i* for a single target is often quantified as a free energy of binding (Δ*G*_*i*,target_), there are a number of different metrics for quantifying compound selectivity [55, 56]. Here, we consider the selectivity *S*_*i*_ between one target and another (an *antitarget*) as the difference in free energy of binding for a given ligand *i* between the two,

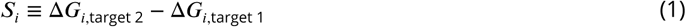

While in the optimization of potency we are concerned with ΔΔ*G*_*ij*_ = Δ.*G*_*j*_ − Δ*G*_*i*_, the relative free energy of binding of ligands *i* and *j* to a single target, in the optimization of selectivity, we are concerned with Δ*S*_*ij*_ ≡ *S*_*j*_ − *S*_*i*_, which reflects the change in selectivity between ligand *i* and a related ligand *j*,

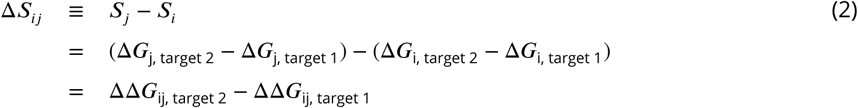

To predict the change in selectivity, Δ*S*_*ij*_, between two related compounds, we developed a protocol that uses a relative free energy calculation (FEP+) [12] to construct a map of alchemical perturbations between ligands in a congeneric series, as described in detail in the **Methods**. The calculation is repeated for each target of interest, with identical perturbations (edges) between each ligand (nodes). Each edge represents a relative alchemical free energy calculation that quantifies the ΔΔ*G* between the ligands (nodes) for a single target. From these calculations, we can then compute the change in selectivity between the two targets of interest, Δ*S*_*ij*_, achieved by transforming ligand *i* into ligand *j*.

Previous work has demonstrated that FEP+ can achieve an accuracy (*σ*_target_) of roughly 1 kcal/mol in single-target potency prediction, which reflects a combination of systematic error and random statistical error [12]. However, it is possible that the systematic error for a given perturbation between ligands *i* and *j* (*σ*_sys, ij, target_) in two different systems may fortuitously cancel when computing Δ*S*_*ij*_, resulting in the systematic contribution to the selectivity error (*σ*_selectivity_) being significantly lower than its contribution to single-target potency error (*σ*_target_). This systematic error may cancel between the two systems for a variety of reasons. For example, a ligand force field parameter assignment error might lead to an spuriously large solvation free energy for a particular compound, which will cancel in the selectivity analysis. Likewise, a sparingly soluble compound might have a similar experimental measurement error for the on-target protein as the off-target protein. Similar cancellation of systematic errors might be observed in ligand and/or protein protonation state assignment error, or systematic differences existing between the protein constructs used for crystallographic studies and biochemical or biophysical assays.

If we presume that the systematic errors for both targets are distributed according to a bivariate normal distribution with correlation coefficient *ρ* quantifying the *degree* of correlation (with *ρ* = 0 denoting no correlation, *ρ* = 1 denoting perfect correlation, and *ρ* = −1 denoting perfect anti-correlation), and that the statistical errors for both targets (*σ*_stat,ij,target_) are completely independent because the simulations for each target are separate, we can model the error in predicting the Δ*S*_*ij*_ as *σ*_selectivity_,

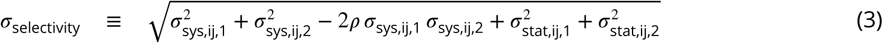

*σ*_selectivity_ can be split into two components: systematic error and statistical error. As more effort is spent on sampling, the per-target statistical error for a given transformation from ligand *i* to ligand *j* (*σ*_stat,ij, target_) will decrease, eventually becoming zero in the regime of infinite sampling. The correlation coefficient *ρ* can be both negative and positive. When the correlation coefficient *ρ* is positive, the systematic error (*σ*_sys, ij, target_) should cancel out, making *σ*_selectivity_ smaller than expected. When the correlation coefficient *ρ* is negative, the systematic error (*σ*_sys, ij, target_) will be anti-correlated, making the *σ*_selectivity_ larger than expected. As we shall see below, the quantitative value of the correlation coefficient *ρ* for the systematic error component has important ramifications for the accuracy with which selectivity can be predicted.

### Correlation in systematic errors can significantly enhance accuracy of selectivity predictions

To demonstrate the potential impact the correlation coefficient *ρ* has on predicting selectivity using alchemical free energy techniques, we created a simple numerical model following Equation 3 which takes into account each of the per-target systematic errors (*σ*_sys,ij,1_, *σ*_sys,ij,2_) expected from the methodology as well as the correlation in those errors, while assuming infinite effort is spent on sampling to reduce the statistical error component (*σ*_stat_) to zero. As seen in Figure 1A, if the per target systematic errors are the same magnitude (*σ*_sys,ij,1_ = *σ*_sys,ij,2_), *σ*_selectivity_ approaches 0 as the correlation coefficient *ρ* approaches 1, even though the single-target potency systematic error is nonzero. If the error for the free energy method is not the same magnitude (*σ*_sys,ij,1_ ≠ *σ*_sys,ij,2_), *σ*_selectivity_ gets smaller but approaches a non-zero value as *ρ* approaches 1.

**Figure 1.**
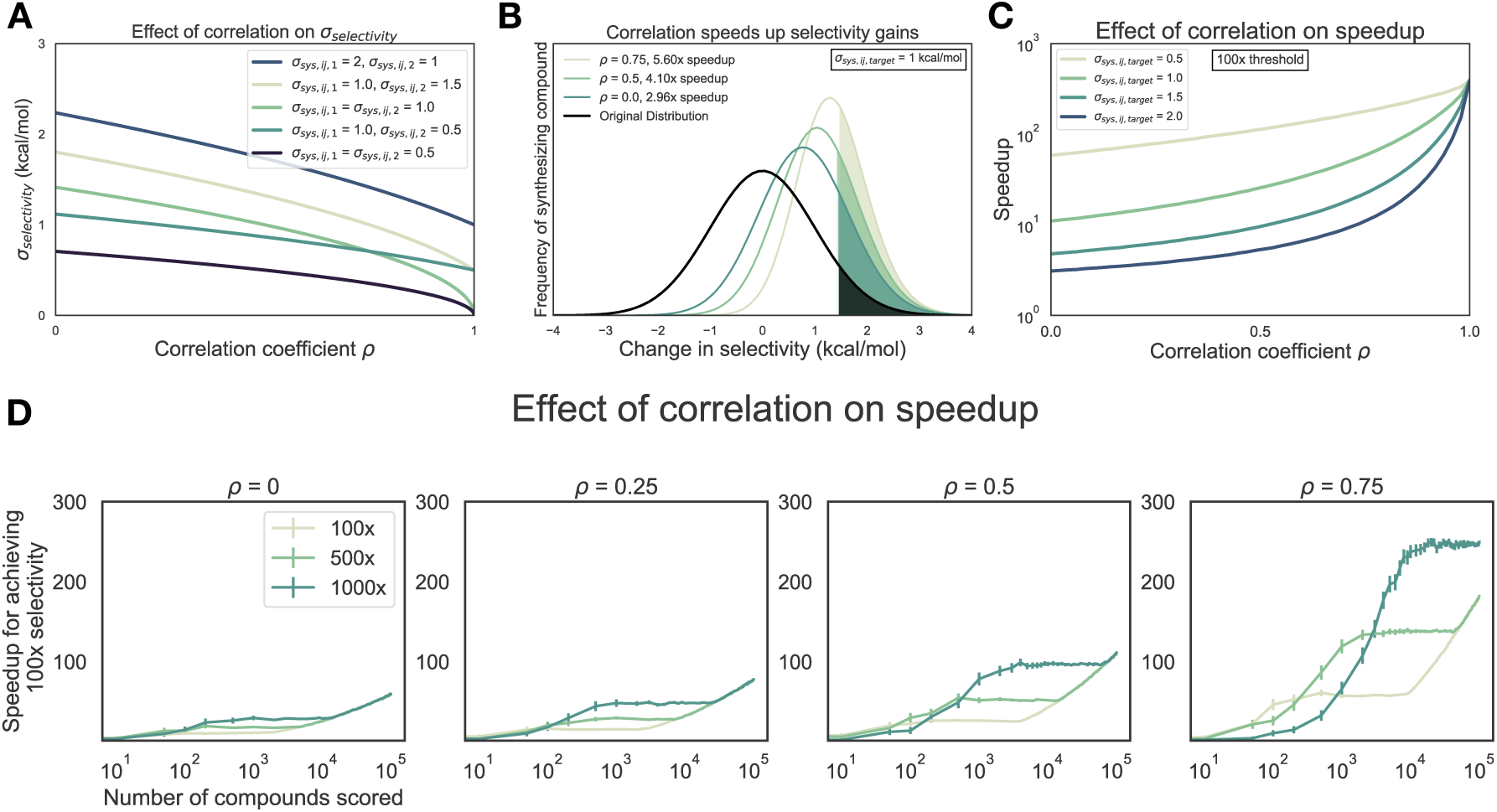
Free energy calculations can accelerate selectivity optimization. (**A**) The effect of correlation on expected errors for predicting selectivity (*σ*_selectivity_) in kcal/mol when statistical error is negligible due to infinite sampling. Each curve represents a different combination of per target systematic errors (*σ*_sys,ij,1_ and *σ*_sys,ij,2_). (**B**) The change in selectivity for molecules proposed by medicinal chemists optimizing a lead candidate can be modeled by a normal distribution centered on zero with a standard deviation of 1 kcal/mol (black curve). Each green curve corresponds to the distribution of compounds made after screening for a 1 log_10_ unit (1.4 kcal/mol) improvement in selectivity with a free energy methodology with a 1 kcal/mol per target systematic error and a particular correlation, in the regime of infinite sampling where statistical error is zero. The shaded region of each curve corresponds to the compounds with a real 1 log_10_ unit improvement in selectivity. The speedup reflects the expected reduction in compounds that must be synthesized to reach a selectivity goal, and is calculated as the ratio of the percentage of compounds made with a real 1 log_10_ unit improvement to the percentage of compounds that would be expected in the original distribution. (**C**) The speedup (y-axis, log scale) expected for 100× (2 log_10_ units, or 2.8 kcal/mol) selectivity optimization as a function of correlation coefficient *ρ*. Each curve corresponds to a different value of *σ*_sys,ij,target_. (**D**) The speedup (y-axis) expected for 100× (2 log_10_ units, or 2.8 kcal/mol) selectivity optimization as a function of number of compounds scored computationally (x-axis) and correlation coefficient *ρ* (each panel) for a method with per-target systematic error (*σ*_sys,ij,target_) of 1 kcal/mol in the regime of infinite sampling. After profiling, the top compounds that meet or surpass the synthesis rule (the predicted selectivity threshold a compound must reach to be triggered for synthesis, each curve) are synthesized, up to a maximum of 10 synthesized compounds. Error bars (y-axis) represent the 95% CI for 1000 replicates at each point. The expected speedup is calculated as the ratio of the number of synthesized compounds that have a true selectivity improvement of 2.8 kcal/mol (100× or 2 log units) divided by the expectation of a compound showing a true selectivity improvement of 2.8 kcal/mol had the same number of compounds that were synthesized been drawn randomly from the underlying unit normal distribution. If no compounds were predicted to meet or surpass the synthesis rule, the speedup was assigned a default value of 1.

To quantify the expected reduction in number of compounds that must be synthesized to achieve a desired selectivity threshold (hereafter referred to as the *speedup* in selectivity optimization), we modeled the change in selectivity with respect to a reference compound for a number of compounds a medicinal chemist might suggest as a normal distribution centered around 0 with a standard deviation of 1 kcal/mol (Figure 1B, black curve), reflecting the notion that most proposed modifications would not drive large changes in selectivity. This assumption—that a synthetic chemist’s proposal distribution can be modeled as a normal distribution—is based on data-driven estimates from an Abbott Laboratories analysis of potency changes [57]

Further suppose that each compound is evaluated computationally with a free energy methodology that has a per-target systematic error (*σ*_sys, ij,target_) of 1 kcal/mol, where we presume sufficient computational effort has been expended to make statistical error negligible. All compounds predicted to have a 1.4 kcal/mol or greater improvement in selectivity (10× in ratio of affinities, or 1 log_10_ unit) are synthesized and experimentally tested (Figure 1B, colored curves), using an experimental technique with perfect measurement accuracy. The fold-change in the proportion of compounds that are made that have a true 1.4 kcal/mol improvement in selectivity compared to the original distribution can be calculated as a surrogate for the expected speedup. For this 1.4 kcal/mol selectivity improvement threshold, a correlation coefficient *ρ* = 0.5 gives an expected speedup of 4.1×, which can be interpreted as needing to make 4.1x fewer compounds to achieve a tenfold improvement in selectivity. This process can be extended for the even more difficult proposition of achieving a hundredfold improvement in selectivity (Figure 1C), where 200–300× speedups can be expected, depending on the single-target systematic error (*σ*_sys,ij,target_) for the free energy methodology.

These estimates represent an ideal scenario, where the number of compounds scored and synthesized is unlimited. In a more realistic discovery project, the number of compounds scored is limited by computational resources, and the number of compounds synthesized is limited by chemistry resources. In this case, the observed speedup will depend not only on the correlation coefficient *ρ* and per-target systematic error (*σ*_sys,ij,target_), but also the number of compounds scored and the synthesis rule, defined as the selectivity threshold a compound must be predicted to reach before being selected for synthesis. To model this process, suppose a given number of compounds (Figure 1D, x-axis of each panel) are profiled with a free energy method with a per-target systematic error (*σ*_sys,ij, target_) of 1 kcal/mol and some correlation coefficient (*ρ*). The top compounds that are predicted to have an improvement in selectivity greater than a set “synthesis rule” threshold (100×, 500×, or 1000×, Figure 1D, each curve) are synthesized, up to a maximum of 10 compounds. The expected speedup can then be calculated as the ratio of the number of synthesized compounds that have a true selectivity improvement of 2.8 kcal/mol (100× or 2 log units) to the number of compounds expected to have a true selectivity improvement of 2.8 kcal/mol had the same number of compounds as were synthesized been drawn randomly from the underlying unit normal distribution.

As shown in Figure 1D, the more stringent synthesis rules combined with high correlation coefficients (*ρ*) allow free energy calculations to have the highest impact in designing selectivity inhibitors, provided enough compounds have been scored. Interestingly, at correlation coefficient *ρ*=0.75 and low numbers of scored compounds, the 500× synthesis provides a greater speedup than 1000× synthesis rule. This is because there is high probability no compounds meet the more 1000× stringent synthesis rule until many more compounds are scored. This has implications for drug discovery efforts, where time and computational effort may limit the number of compounds able to be profiled with free energy methods.

### An experimental data set of CDK2/CDK9 inhibitors demonstrates the difficulty in achieving high selectivity

To assess the correlation of errors in free energy predictions for selectivity, we set out to gather data sets that met a number of criteria. We searched for data sets that contained binding affinity data for a number of kinase targets and ligands in addition to crystal structures for each target with the same ligand.

This data set contains a congeneric series of ligands with experimental data for CDK2 and CDK9, with the goal of potently inhibiting CDK9 and sparing CDK2. Based on a multiple sequence alignment of the 85 binding site residues identified in the kinase–ligand interaction fingerprints and structure (KLIFS) database [58, 59], CDK2 and CDK9 share 57% sequence identity (Table S1, Figure S1). For this CDK2/CDK9 data set [53], ligand 12c was cocrystallized with CDK2/cylin A (Figure 2A, left) and CDK9/cyclin T (Figure 2B, left), work that was published in a companion paper [60]. In both CDK2 and CDK9, ligand 12c forms relatively few hydrogen bond interactions with the kinase. Each kinase forms a pair of hydrogen bonds between the ligand scaffold and a hinge residue (C106 in CDK9 and L83 in CDK2) that is conserved across all of the ligands in this series. CDK9, which has slightly lower affinity for ligand 12c (Figure 2C, right), forms an interaction between the sulfonamide of ligand 12c and residue E107. On the other hand, CDK2 forms interactions between the sulfonamide of ligand 12c and residues K89 and H84. The congeneric series of ligands contains a number of difficult perturbations, particularly at substituent point R3 (Figure 2C, left). Ligand 12i also presented a challenging perturbation, moving the 1-(piperazine-1-yl)ethanone from the *meta* to *para* location.

**Figure 2.**
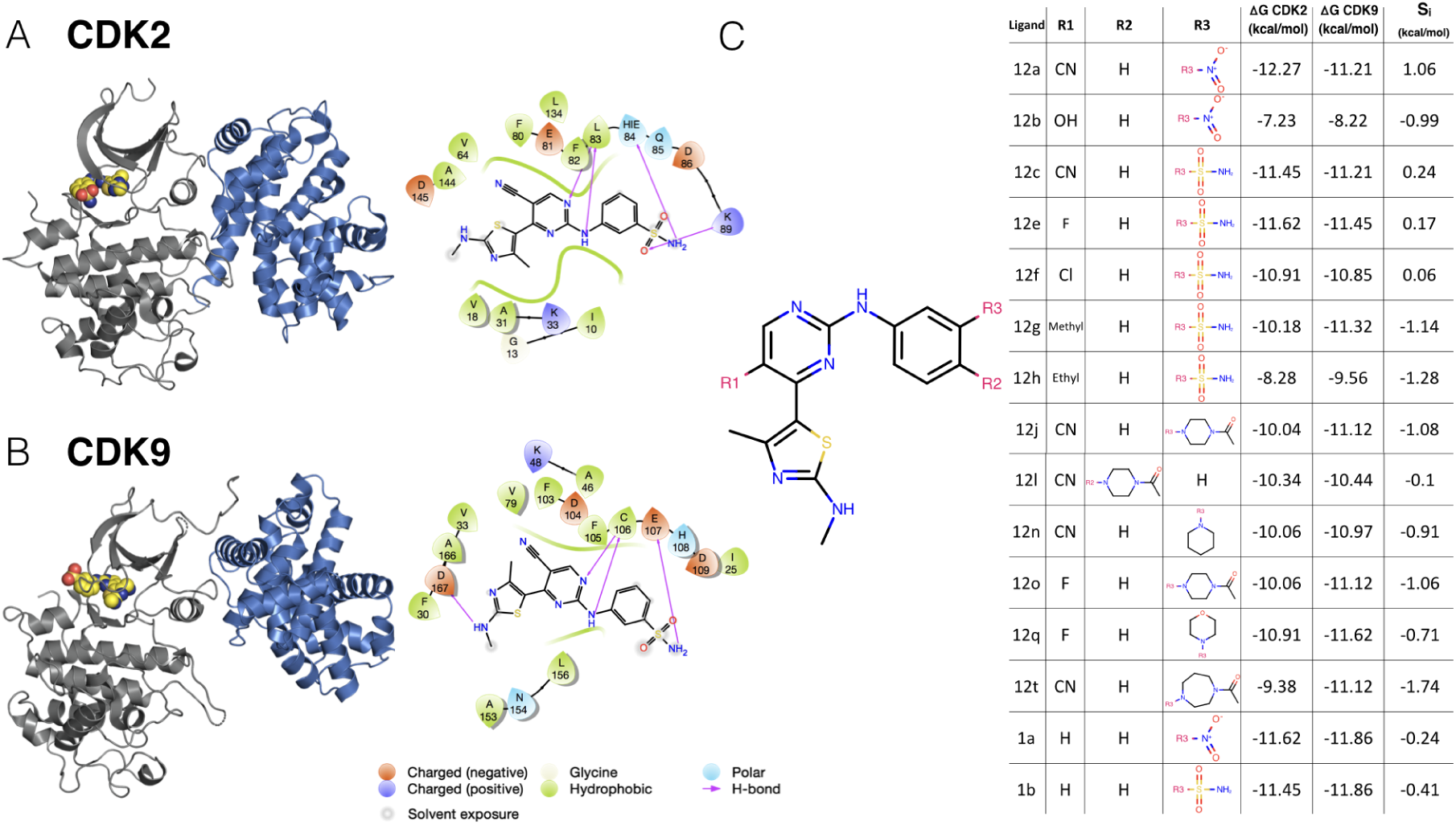
A CDK2/CDK9 data set illustrates selectivity optimization between closely-related kinases. Experimental IC_50_ data for a congeneric series of compounds binding to CDK2 and CDK9 was extracted from Shao et al. [53] and converted to free energies of binding. (**A**) *(left)* Crystal Structure (4BCK) [60] of CDK2 (gray ribbon) bound to ligand 12c (yellow spheres). Cyclin A is shown in blue ribbon. *(right)* 2D ligand interaction map of ligand 12c in the CDK2 binding site. (**B**) *(left)* Crystal structure of CDK9 (4BCI)[60] (gray ribbon) bound to ligand 12c (yellow spheres). Cyclin T is shown in blue ribbon. *(right)* 2D ligand interaction map of ligand 12c in the CDK9 binding site. (**C**) *(left)* 2D structure of the common scaffold for all ligands in congeneric ligand series 12 from the publication. *(right)* A table summarizing all R group substitutions as well as the published experimental binding affinities and selectivities [53], derived from the reported *K*_*i*_ as described in **Methods**.

This congeneric series of ligands also highlights two of the challenges of working from publicly available data: First, the dynamic range of selectivity is incredibly narrow, with a mean *S* (CDK9 - CDK2) of - 0.65 kcal/mol, and a standard deviation of only 0.88 kcal/mol; the total dynamic range of this data set is 2.8 kcal/mol. Second, experimental uncertainties are not reported for the experimental measurements. This data set reported *K*_*i*_ values calculated from measured IC_50_, using the *K*_*m*_ (ATP) for CDK2 and CDK9 and [ATP] from the assay using the Cheng-Prussof equations [61]. Thus, for this and subsequent sets of ligands, the random experimental uncertainty is assumed to be 0.3 kcal/mol based on previous work done to summarize uncertainty in experimental data, assuming there is no systematic experimental error. While *K*_*i*_ values are reported, these values are derived from IC50 measurements. A number of studies report on the reproducibility of intra-lab IC50 measurements. These values range from as low as 0.22 kcal/mol [62], from public data, to as high as 0.4 kcal/mol [6], which was estimated from internal data at Abbott Laboratories. The assumed value of 0.3 kcal/mol falls within this range, and agrees well with the uncertainty reported from Novartis for two different ligand series [63].

### An experimental data set of CDK2/ERK2 inhibitors where greater selectivity was achieved

The CDK2/ERK2 data set from Blake et al. [54] also met the criteria described above, with the goal of developing a potent ERK2 inhibitor. Based on a multiple sequence alignment of the KLIFs binding site residues [58, 59], CDK2 and ERK2 share 52% sequence identity (Table S1, Figure S1), making them slightly less closely related than CDK2 and CDK9 (57%). Note that while all three kinases belong to the CMGC family and are closely related in the phylogenetic Manning tree, CDK2 and CDK9 belong to the CDK (Cyclin-dependent kinase) subfamily, while ERK2 is part of the nearby MAPK (Mitogen-activated protein kinases) subfamily. From a structural point of view, the two kinase pdb pairs used in this study are also very similar. Binding site superposition revealed that both pdb pairs align well, only a marginally lower RMSD of 0.81 A was obtained for the CDK2/CDK9 pair compared to 0.92 A for CDK2/ERK2 pair.

Crystal structures for both CDK2 (Figure 3A, top) and ERK2 (Figure 3B, top) were available with ligand 22 (according to the manuscript numbering scheme) co-crystallized. Of note, CDK2 was not crystallized with cyclin A, despite cyclin A being included in the affinity assay reported in the paper [54].

**Figure 3.**
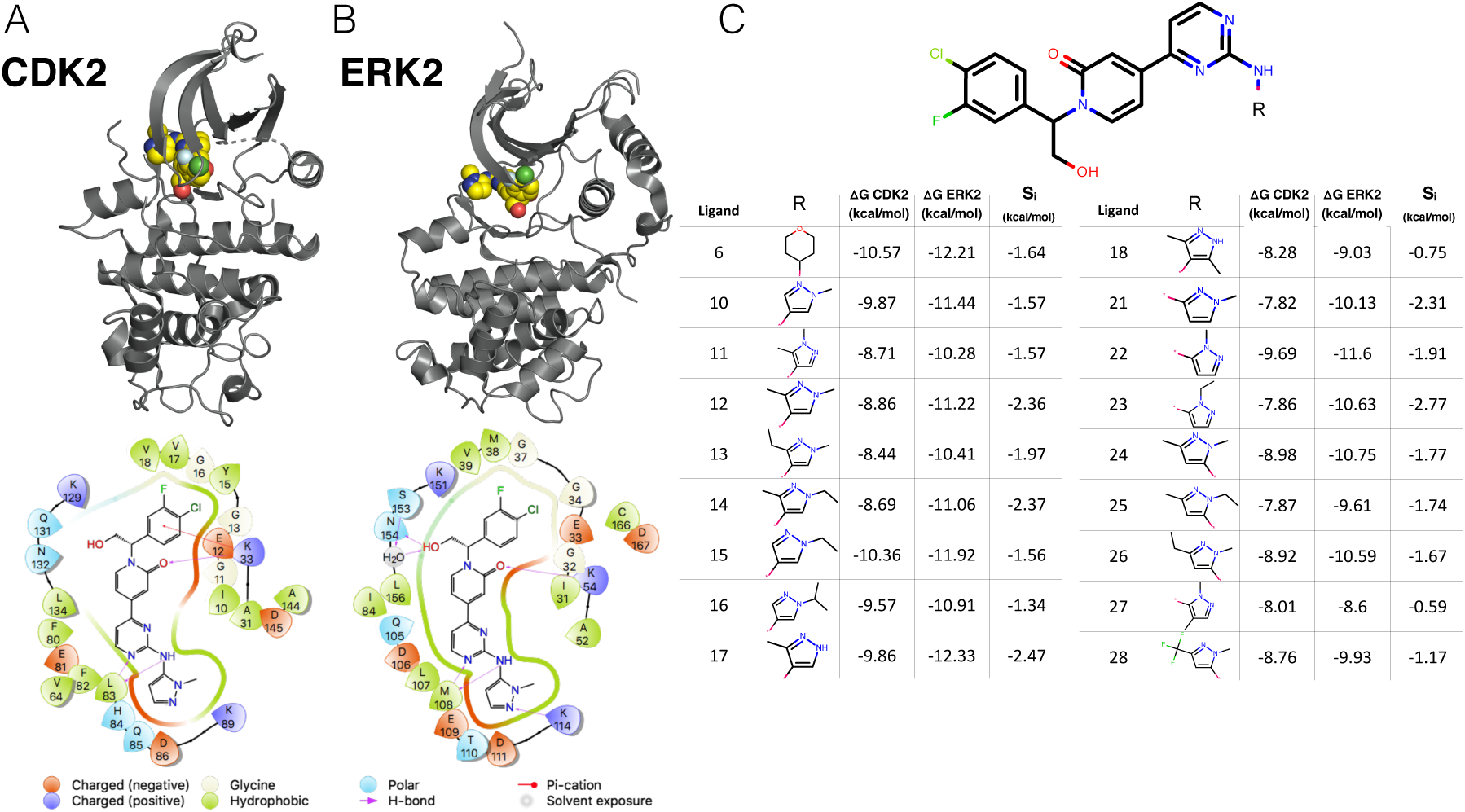
A CDK2/ERK2 data set illustrates selectivity optimization among more distantly related kinases. (**A**) *(top)* Crystal structure of CDK2 (5K4J) shown in gray cartoon and ligand 22 shown in yellow spheres. *(bottom)* 2D interaction map of ligand 22 in the binding pocket of CDK2 (**B**) *(top)* Crystal structure of ERK2 (5K4I) shown in gray cartoon with ligand 22 shown in yellow spheres. *(bottom)* 2D interaction map of ligand 22 in the binding pocket of ERK2. (**C**) *(top)* Common scaffold for all of the ligands in the Blake data set [54], with R denoting attachment side for substitutions. *(bottom)* Table showing R group substitutions and experimentally measured binding affinities and selectivities, derived from the IC_50_ values as described in the methods section. Ligand numbers correspond to those used in the Blake publication [54].

CDK2 in this crystal structure (4BCK) adopts a DFG-in conformation with the *α*C helix rotated out, away from the ATP binding site and breaking the conserved salt bridge between K33 and E51 (Figure S2A), indicative of an inactive kinase [44, 64]. By comparison, the CDK2 structure from the CDK2/CDK9 data set adopts a DFG-in conformation with the *α*C helix rotated in, forming the ionic bond between K33 and E51 indicative of an active kinase, due to allosteric activation by cyclin A. While missing cyclins have caused problems for free energy calculations in prior work, it is possible that the fully active, cyclin-bound conformation contributes equally to binding affinity for all of the ligands in this series, and the high accuracy of the potency predictions (Figure 4, top left) is the result of fortuitous cancellation of errors.

**Figure 4.**
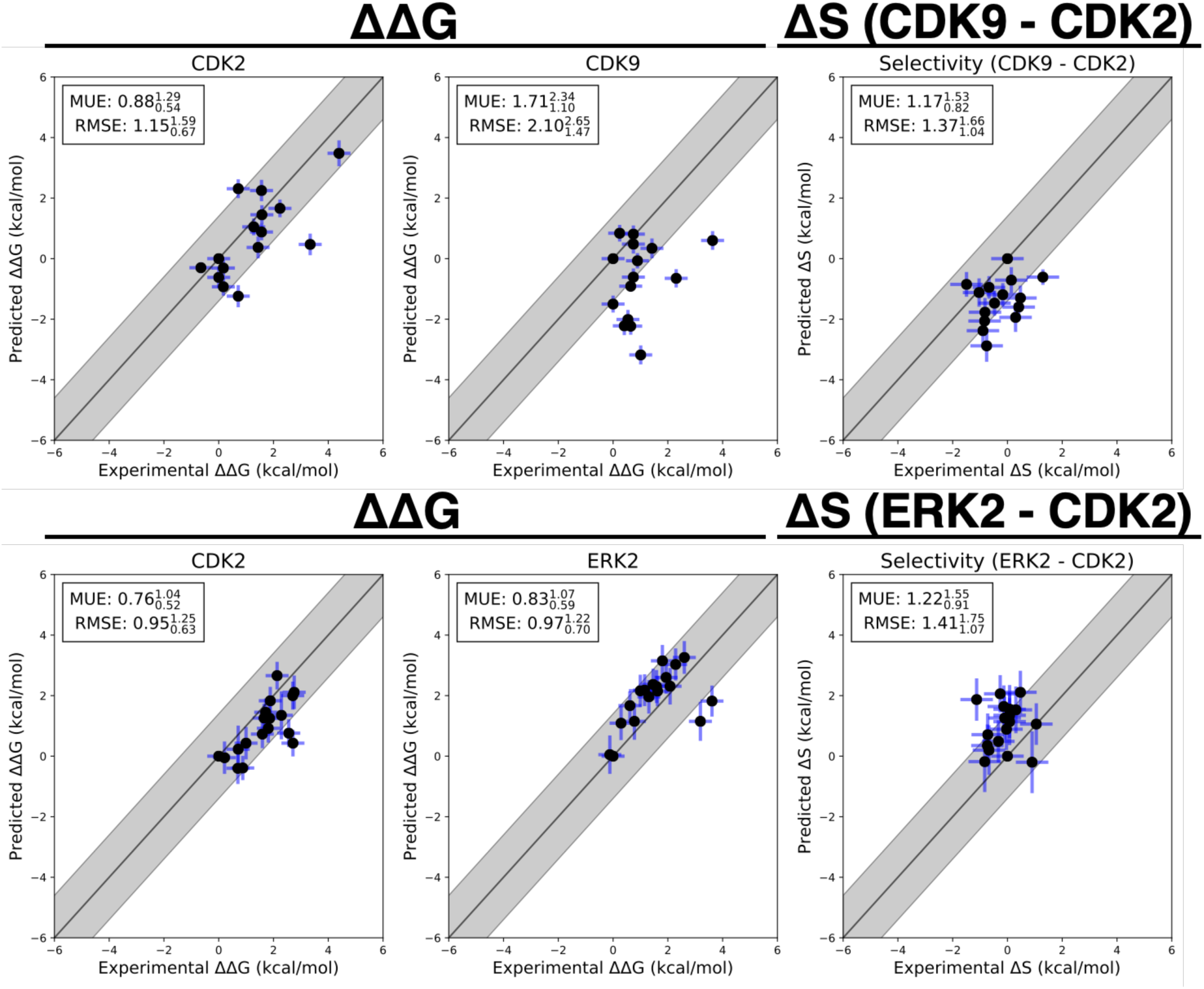
Selectivity predictions suggest correlation in systematic error. Δ Δ*G*_*i*,ref,target_ and Δ*S*_*i*,ref_ predictions for CDK2/CDK9 (*top*) from the Shao data sets and CDK2/ERK2 from the Blake data sets (*bottom*). The experimental values are shown on the X-axis and calculated values on the Y-axis. Each data point corresponds to a transformation between a ligand *i* to a set reference ligand (ref) for a given target. All values are shown in units of kcal/mol. The horizontal error bars show to the 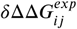 based on the assumed uncertainty of 0.3 kcal/mol[6, 63] for each 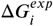. We show the estimated statistical error (*σ*stat,ij,target) as vertical blue error bars, which are one standard error. For selectivity, the errors were propagated under the assumption that they were completely uncorrelated. *σ*_stat,ij,target_ was estimated by calculating the standard deviation of 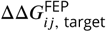 from the Bayesian model described in depth in **Methods**, where *j* is the reference compound. The black line indicates agreement between calculation and experiment, while the gray shaded region represent 1.36 kcal/mol (or 1 log_10_ unit) error. The mean unsigned error (MUE) and root-mean squared error (RMSE) are shown on each plot with bootstrapped 95% confidence intervals.

The binding mode for this series is similar between both kinases. There is a set of conserved hydrogen bonds between the scaffold of the ligand and the backbone of one of the hinge residues (L83 for CDK2 and M108 for ERK2). The conserved lysine (K33 for CDK2 and K54 for ERK2), normally involved in the formation of a ionic bond with the *α*C helix, forms a hydrogen bond with the scaffold (Figure 3A and 3B, bottom) in both CDK2 and ERK2. However, in the ERK2 structure, the hydroxyl engages a crystallographic water as well as N154 in a hydrogen bond network that is not present in the CDK2 structure. The congeneric ligand series features a single solvent-exposed substituent. This helps to explain the narrow distribution of selectivities, with a mean selectivity of −1.74 kcal/mol (ERK2 - CDK2) and standard deviation of 0.56 kcal/mol; the total dynamic range of this data set is 2.2 kcal/mol. While the small standard deviation suggests that selectivity is difficult to drive with R-group substitution, the total dynamic range demonstrates that R-group substitutions can provide significant selectivity enhancements.

### FEP+ calculations show smaller than expected errors for CDK2/CDK9 Δ*S*_*ij*_ predictions

Three replicates of FEP+ calculations were run on each target for both experimental data sets described above. The FEP+ predictions of the relative free energy of binding between ligands *i* and a reference compound (ref) for each target (Δ Δ*G*_*i*,ref,*target*_) showed good accuracy and consistent results for all three replicates. The results for replicate 1 are reported in Figure 4 for both the CDK2 and ERK2 data set (bottom) and the CDK2/CDK9 data set (top), Δ Δ*G*_*i,ref*,target_ is defined for each ligand *i* using a consistent reference compound within data sets.

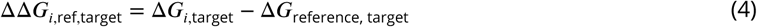

The reference compounds (Compound 6 for CDK2/ERK2 and Compound 1a for CDK2/CDK9) were selected because they were the initial compounds from which the reported synthetic studies were started. Replicate 1 of the CDK2/ERK2 calculations is shown on the bottom of Figure 4, with an RMSE of 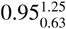 and 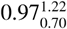 kcal/mol to CDK2 and ERK2, respectively (where the lower and upper values indicate a 95% confidence interval). The RMSE reported here is calculated for all of the Δ Δ*G*_*i,ref*,target_ that were predicted. All of the CDK2 and ERK2 Δ Δ*G*_*i,ref*,target_s were predicted within 1 log unit of the experimental value. The change in selectivity (Δ*S*_*ij*_) predictions show an RMSE of 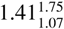 kcal/mol, with all the confidence intervals of the predictions falling within 1 log unit of the experimental values (Figure 4, top right panel). This RMSE is comparable to the expected RMSE of 1.36, assuming the error from the CDK2/ERK2 calculations behaves in an uncorrelated manner (Equation 3 where the correlation coefficient *ρ* is zero). This was consistent across all three replicates of the calculations (Figure S6).The narrow dynamic range for selectivity combined with high experimental and computational uncertainty highlight the challenges for predicting selectivity. When the error of the calculated selectivity is comparable to the dynamic range of selectivity, then the calculations cannot predict with statistical confidence whether any compound is more selective than the other.

Replicate 1 of the CDK2/CDK9 calculations are shown in the top panel of Figure 4. The CDK2 and CDK9 data sets show higher errors in Δ Δ*G*_*i*,ref,target_ predictions, with an RMSE of 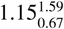 and 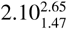 kcal/mol respectively. This higher RMSE is driven by the reference compound, (Compound 1a) being poorly predicted, particularly in CDK9. There are a number of outliers that fall outside of 1 log_10_ unit from the experimental value for CDK9. While the higher per target errors make predicting potency more difficult, the selectivity predictions show an RMSE of 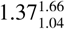 kcal/mol. This observed RMSE is lower than what would be expected if the error were completely uncorrelated between CDK2 and CDK9, propagated as in Equation 3 where the correlation coefficient *ρ* is zero to get an expected value of 2.38 kcal/mol. This suggests that some correlation in the error is leading to fortuitous cancellation of the systematic error, leading to more accurate than expected predictions of Δ*S*_*ij*_. These results were consistent across all three replicates of the calculation (Figure S4).

### Correlation of systematic errors accelerates selectivity optimization

To quantify the correlation coefficient (*ρ*) of the systematic error between targets, we built a Bayesian graphical model to separate the systematic error from the statistical error and quantify our confidence in estimates of *ρ* (described in depth in Methods). Briefly, we modeled the absolute free energy (*G*) of each ligand in each thermodynamic phase (ligand-in-complex and ligand-in-solvent, with *G* determined up to an arbitrary additive constant for each phase) as in Equation 15. The model was chained to the FEP+ calculations by providing the 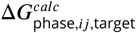 for each edge from the FEP+ maps (where *j* is now not necessarily the reference compound) as observed data, as in Equation 17. As in Equation 19, the experimental data was modeled as a normal distribution centered around the true free energy of binding 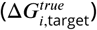 corrupted by experimental error, which is assumed to be 0.3 kcal/mol from previous work done to quantify the uncertainty in publicly available data [6]. Δ*G* values derived from reported IC_50_s or *K*_*i*_s, as described in the methods section, were treated as data observations (Equation 19) and the 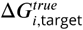 was assigned a weak normal prior (Equation 20).

The correlation coefficient *ρ* was calculated for each Bayesian sample from the model posterior according to equation 22. The CDK2/CDK9 calculations show strong evidence of correlation, with a correlation coefficient of 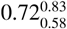 (Figure 5A, right) for replicate 1. The rest of the replicates showed strong agreement (Figure S4). The joint marginal distribution of the error (*ϵ*) for each target (Figure 5A, left) is more diagonal than symmetric, which is expected for cases in which *ρ* is high (Figure S3).

**Figure 5.**
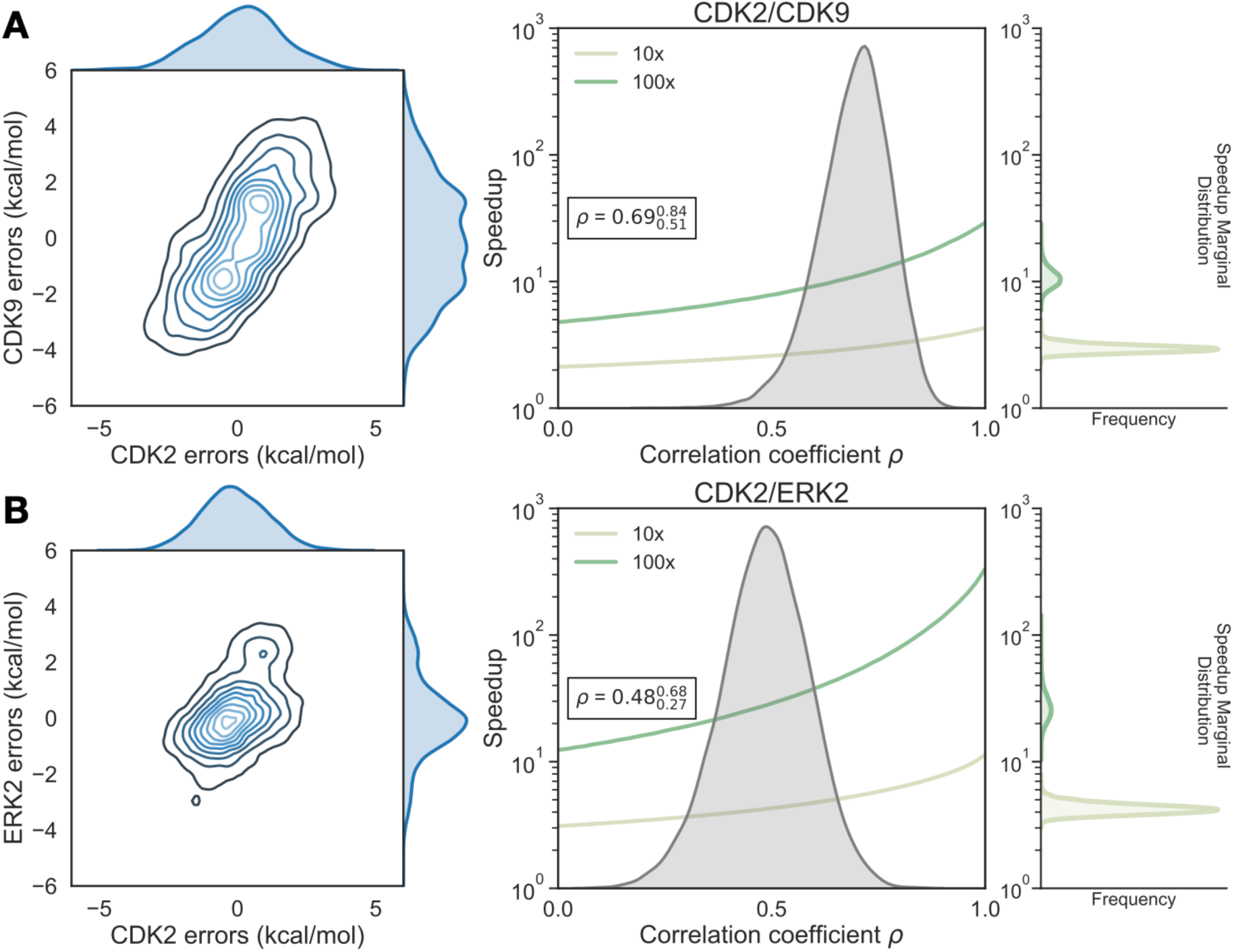
Correlation in systematic errors between targets can significantly accelerate selectivity optimization. (**A**, *left*) The joint posterior distribution of the prediction errors for the more distantly related CDK2 (x-axis) and CDK9 (y-axis) from the Bayesian graphical model. (**A**, *right*) Speedup in selectivity optimization (y-axis), which estimates the reduction in compounds that must be synthesized to achieve a target selectivity when aided by free energy calculations, using the model where the number of compounds scored and synthesized is unlimited, as a function of correlation coefficient (x-axis). To calculate *σ*_selectivity_, we calculate the per target systematic error (*σ*_sys,ij,target_) by taking the mean of *ϵ*_*ij*,target_ where *j* is the reference compound 1a. The posterior marginal distribution of the correlation coefficient (*p*) is shown in gray, while the expected speedup is shown for 100× (green curve) and 10× (yellow curve) selectivity optimization. The inserted box shows the mean and 95% confidence interval for the correlation coefficient. The marginal distribution of speedup is shown on the right side of the plot for both 100× (green) and 10× (yellow) selectivity optimization speedups. (**B**) As above, but for the more closely related CDK2/ERK2 selectivity data set using compound 6 as the reference.

To quantify the expected speedup of selectivity with this level of correlation in the systematic errors for CDK2/CDK9, we first calculated the per target systematic error *σ*_sys,ij,target_ by taking the mean of the absolute value of *ϵ*_*ij*,target_ where *j* is the reference compound 1a. Combining these estimates for the correlation coefficient (*ρ*) and the per target systematic errors (*σ*_sys,ij,target_), we can compute *σ*_selectivity_ and the expected speedup in the regime of infinite sampling effort where there is no statistical error when the number of compounds scored and synthesized is unlimited. The high correlation in errors for the CDK2/CDK9 calculations leads to a speedup of 3x for 1 log_10_ unit selectivity optimization and 10x for 2 log_10_ unit selectivity optimization (Figure 5A, right), despite the much high per target systematic errors (*σ*_sys,ij,target_).

The correlation coefficient *ρ* for replicate 1 of the CDK2/ERK2 calculations was quantified to be 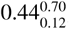, indicating that the errors are moderately correlated between ERK2 and CDK2 (Figure 5B, right); this was consistent with the distribution for *ρ* in replicate 3 (Figure S7), while the confidence interval of *ρ* for replicate 2 is much wider, indicating the correlation is weak.

Considering the speedup model where the number of compounds scored and synthesized is unlimited, the modest correlation and low per target systematic errors for the CDK2/ERK2 calculations allow for a predicted 4–5x speedup for 1 log_10_ unit selectivity optimization, and a 30–40x speedup for 2 log_10_ unit selectivity optimization (Figure 5B, right).

Using the correlation coefficient (*ρ*), *σ*_stat,ij,target_, and *σ*_sys,ij,target_ quantified from the Bayesian model for each set of calculations, we can now calculate the y-axis error bars for the Δ*S* panels of Figure 4 according to the proposed *σ*_selectivity_ equation (Eq 3). Shown in Figure S9, we can see that *σ*_selectivity_ accounts for most of the disagreement between the predicted Δ*S*_*ij*_ and the experimental Δ*S*_*ij*_.

### Expending more effort to reduce statistical error can be beneficial in selectivity optimization

Up to this point, we have considered only systematic error in quantifying the speedup free energy calculations can enable for selectivity optimization, by assuming enough sampling is done to reduce the statistical error for each target to zero. To begin understanding how statistical error impacts this speedup, we modified the model of speedup by additionally considering the per target statistical error (*σ*_stat, target_), which we define in Equation 7 such that at the baseline effort, *N, σ*_stat,ij,target_ is 0.2 kcal/mol. In this definition, it takes 4× the sampling, or effort, to reduce statistical error by a factor of 2×. We assume that statistical error is uncorrelated when propagating to two targets, and that *σ*_sys,ij,target_ is ≈ 1.0 kcal/mol for both targets [4, 62]. As shown in Figure 6, expending effort to reduce *σ*_stat,ij,target_ when *ρ* is less than 0.5 does not change the expected speedup for the 100× selectivity threshold in meaningful way, suggesting that it is not worth running calculations longer than the default protocol in this case. However, when *ρ >* 0.5, the curves do start to separate, particularly the 1/4×, 1×, and 4× effort curves. This suggests that when the correlation is high, running longer calculations can produce net improvements in selectivity optimization speed. Interestingly, the 16×, 48×, and ∞ effort curves do not differ greatly from the 4× effort curve, indicating that there are diminishing returns to running longer calculations.

**Figure 6.**
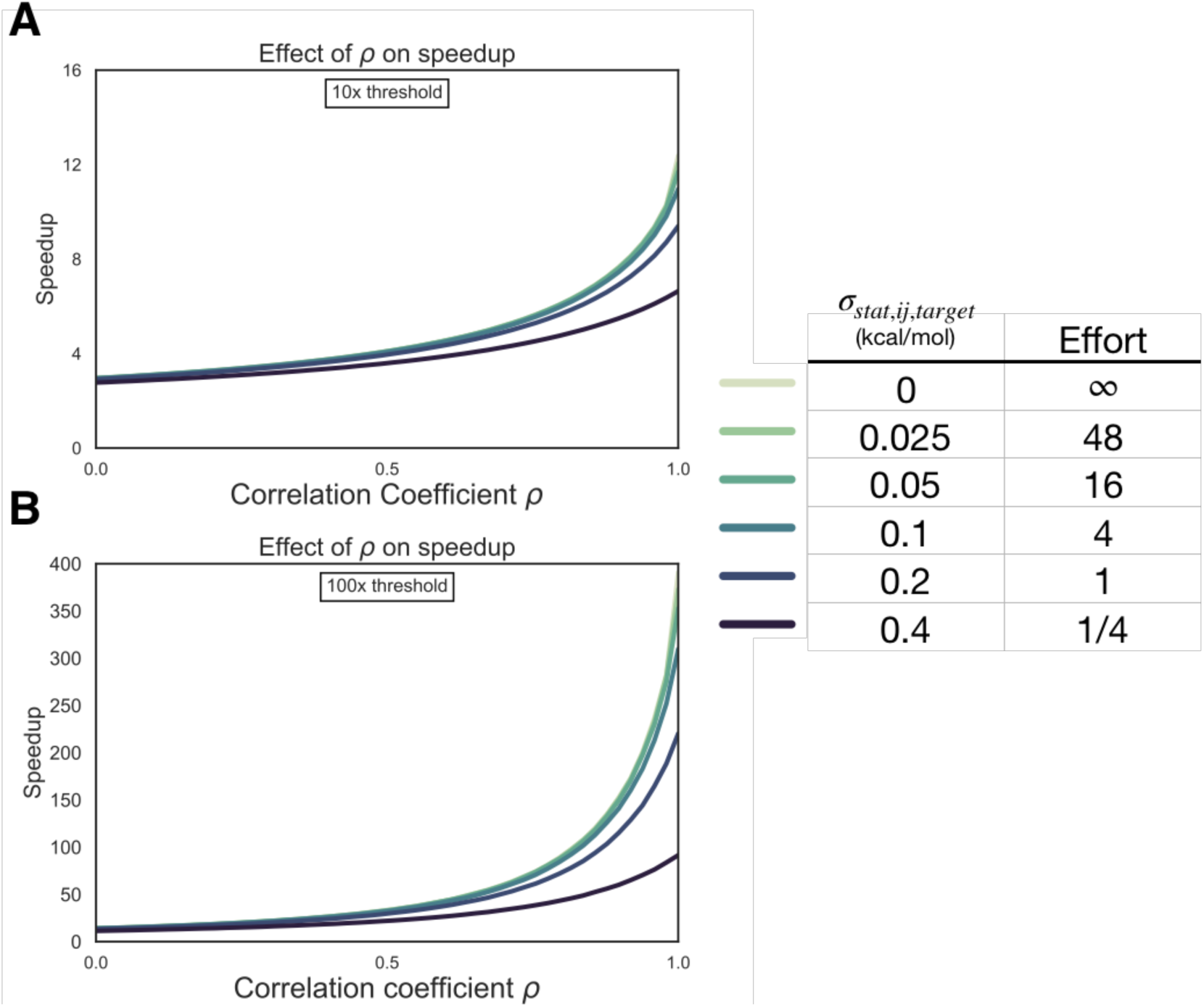
Reducing statistical uncertainty when systematic error correlation is high improves the speedup in selectivity optimization achievable with free energy calculations. (*left*) The speedup in selectivity (Y-axis) as a function of correlation coefficient (X-axis). Each curve represents a different per target statistical error (*σ*_stat,ij,target_) for 10× (1 log_10_ unit) (**A**) and 100× (2 log_10_ unit) (**B**) thresholds (*right*) Table with the per target statistical error (*σ*_stat,ij,target_), kcal/mol) corresponding to each curve on the left and a rough estimate of the generic amount of computational effort it would take to achieve that statistical uncertainty.

### The estimated correlation coefficient is robust to Bayesian model assumptions

In order to better understand the statistical error in our calculations, we performed three replicates of our calculations, and calculated the standard deviation of the cycle closure corrected Δ Δ*G* for each edge of the map, and compared that value to the cycle closure errors and Bennett errors reported for each edge (Figure S8). For each set of calculations, the standard deviation suggests that the statistical error is between 0.1 and 0.3 kcal/mol, which is in good agreement with the reported Bennett error (Figure S8). However, hysteresis in the closed cycles in the FEP map as reflected by the cycle closure error estimates indicate much larger sampling errors than those estimated by the Bennett method or standard deviations of multiple runs, suggesting that both the Bennett errors and standard deviation of multiple replicates are underestimating the statistical error for these calculations. Based on this observation, we include a scaling parameter *α* in the Bayesian error model (Eq. 16) to account for the BAR errors underestimating the cycle closure statistical uncertainty. We also considered using a distribution with heavier tails, such as a Student’s t-distribution, but found the quantification of the correlation coefficient *ρ* insensitive to the use of either a scaling parameter or heavier-tailed distributions (Figure S10).

## Discussion and Conclusions

### *S* is a useful metric for selectivity in lead optimization

There are a number of different metrics for quantifying the selectivity of a compound [55], which look at selectivity from different views depending on the information trying to be conveyed. One of the earliest metrics was the standard selectivity score, which conveyed the number of inhibited kinase targets in a broad scale assay divided by the total number of kinases in the assay [65]. The Gini coefficient is a method that does not rely on any threshold, but is highly sensitive to experimental conditions because it is dependent on percent inhibition [66]. Other metrics take a thermodynamic approach to kinase selectivity and are suitable for smaller panel screens [67, 68]. Here, we propose a more granular, thermodynamic view of selectivity that is straightforward to calculate using free energy methods: the change in free energy of binding for a given ligand between two different targets (*S*). *S* is a useful metric of selectivity in lead optimization once a single, or small panel, of off-targets have been identified and the goal is to use physical modeling to either improve or maintain selectivity within a lead series.

### Systematic error correlation can accelerate selectivity optimization

We have demonstrated, using a simple numerical model that assumes unlimited synthetic and computational resources, the impact that free energy calculations with even weakly correlated systematic errors can have on speeding up the optimization of selectivity in small molecule kinase inhibitors. While the expected speedup is dependent on the per target systematic error of the method (*σ*_sys,ij,target_), the speedup is also highly dependent on the correlation of errors made for both targets. Unsurprisingly, free energy methods have greater impact as the threshold for selectivity optimization goes from 10× to 100×. While 100× selectivity optimization is difficult to achieve, the expected benefit from free energy calculations is also quite high, with speedups of one or two orders of magnitude possible. In a more realistic scenario, where the number of compounds scored and synthesized is limited by resources, we have demonstrated using the same numerical model that more stringent synthesis rules results in increased speedup from free energy calculations. This holds true across different correlation coefficients (*ρ*), provided enough compounds are scored. As our model shows, it is possible for stringent synthesis rules to provide benefits similar to operating with high systematic error correlation coefficients (*ρ*).

### Two pairs of kinase test systems suggest systematic errors can be correlated

To quantify the correlation of errors in two example systems, we gathered experimental data for two congeneric ligand series with experimental data for CDK2 and ERK2, as well as CDK2 and CDK9. These data sets, which had crystal structures for both targets with the same ligand co-crystallized, exemplify the difficulty in predicting selectivity. The dynamic range of selectivity for both systems is very narrow, with most of the perturbations not having a major impact on the overall selectivity achieved. Further, the data was reported without reliable experimental uncertainties, which makes quantifying the errors made by the free energy calculations difficult. This issue is common when considering selectivity, as many kinase-oriented high throughput screens are carried out at a single concentration and not highly quantitative.

The CDK9 calculations contained an outlier, compound 12h, that drove much of the prediction error for that set. Compound 12e (R1 = F) is the most potent against CDK9 of the compounds in with a sulfonamide at R3 (Figure 2). The addition of a single methyl group decreases the potency against CDK9 (compound 12g) and while only slightly changing the affinity for CDK2. However, adding on another methyl group (compound 12h) results in an order of magnitude decrease in *K*_*i*_ for both CDK9 and CDK2. Crystal structures for both kinases show that R1 points into a pocket formed by the backbone, and the sidechains of a Valine and Phenylalanine. While ethyl at R1 in compound 12h *is* bulkier, the magnitude of the decrease in affinity for both kinases is larger than might be expected, given that the pocket suggests an ethyl group would be well accommodated in terms of fit and the hydrophobicity of the sidechains. For both kinases, the free energy calculations predict that this addition should *improve* the potency, suggesting that it is possible that the model is missing a chemical detail that might explain the trend seen in the experimental data. We expect that these types of errors, which would be troubling when predicting potency alone, will drive the correlation of systematic errors and fortuitously cancel when predicting selectivity.

Despite CDK2 and ERK2 belonging to different kinase subfamilies, the calculated correlation in the systematic error for two of the replicates suggests that fortuitous cancellation of errors may be applicable in a wider range of scenarios than closely related kinases within the same subfamily. This may be driven by relatively high binding site sequence identity between CDK2 and ERK2 (52% compared for 57% for CDK2/CDK9). However, the confidence interval of the correlation is quite broad, including 0 for the lower bound for the third replicate, suggesting that errors for more distantly related proteins will have only moderate, if any, correlation.

### Reducing statistical error is beneficial when systematic errors are correlated

In order to better understand if there are situations where it is beneficial to run longer calculations to minimize statistical error to achieve a larger speedup in the synthesis of selective compounds, we built a numerical model of the impact of statistical error in the context of different levels of systematic error correlation. Our results suggest that unless the correlation coefficient *ρ* is highly positive for the two targets of interest, there is not much benefit in running longer calculations. However, when the systematic error is reduced by correlation, longer calculations can help realize large increases in speedup to achieve selectivity goals. Keeping a running quantification of *ρ* for free energy calculations as compounds are made and the predictions can be tested will allow for decisions to be made about whether running longer calculations is worthwhile. It will also allow for an estimate of *σ*_selectivity_, which is useful for estimating expected systematic error for prospective predictions. Importantly, we expect that correlation will be modeling protocol dependent and any changes to the way the system is modeled over the course of discovery program are expected to change the observed correlation in the systematic error.

### Larger data sets with a wide range of protein targets will enable future work

The data sets gathered here were limited by the total number of compounds, the small dynamic range for selectivity (*S*), and the lack of reliable experimental uncertainties. The small size of the data set makes it difficult to draw broad conclusions about the correlation in systematic errors. Understanding the degree of correlation *a priori* based on structural or sequence similarity requires study on a larger range of targets than the two pairs presented in this study. A larger data set that contained many protein targets, crystal structures, and quantitative binding affinity data would be ideal to draw conclusions about the broader prevalence of systematic error correlation.

This work demonstrates that correlation in the systematic errors can allow free energy calculations to facilitate significant speedups in selectivity optimization for drug discovery projects. This is particularly important in kinase systems, where considering multiple targets is an important part of the development process. The results suggest that free energy calculations can be particularly helpful in the design of kinase polypharmacological agents, especially in cases where there is high correlation in the systematic errors between multiple targets.

## Methods

### Numerical model of selectivity optimization speedup

To model the impact correlation of systematic error would have on the expected uncertainty for selectivity predictions, *σ*_selectivity_ was calculated using Equation 3 for 1000 evenly spaced values of the correlation coefficient (*ρ*) from 0 to 1, for a number of combinations of per target systematic errors (*σ*_sys,ij,1_ and *σ*_sys,ij,2_). In the regime of infinite sampling and zero statistical error, the second term reduces to zero.

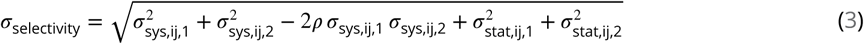

The speedup in selectivity optimization that could be expected from using free energy calculations of a particular per target systematic error (*σ*_sys,ij,target_) was quantified as follows using NumPy (v 1.14.2). An original, true distribution for the change in selectivity of 200 000 000 new compounds proposed with respect to a reference compound was modeled as a normal distribution centered around 0 with a standard deviation of 1 kcal/mol. This assumption was made on the basis that the majority of selectivity is driven by the scaffold, and R group modifications will do little to drive changes in selectivity. The 1 kcal/mol distribution is supported by the standard deviations of the selectivity in the experimental data sets referenced in this work, which are all less than, but close to, 1 kcal/mol.

In this model, we suppose that each of proposed compound is triaged by a free energy calculation and only proposed compounds predicted to increase selectivity by Δ*S*_ij_ ≥1.4 kcal/mol (1 log_10_ unit) with respect to a reference compound would be synthesized. Based on reported estimates in the literature, we presume that relative free energy calculations have a per-target systematic error *σ*_sys,ij,target_ 1 kcal/mol [4], and explore the impact of the correlation coefficient *ρ* governing the correlation of these predictions between targets. The standard error in predicted selectivity, *σ*_selectivity_, is given by Equation 3. When sampling is infinite and *σ*_stat,ij,target_ is zero, *σ*_selectivity_ is driven entirely by the systematic error component (*σ*_sys,ij,target_), resulting in the error in predicted change in selectivity Δ*S*_*ij*_ modeled as a normal distribution centered around 0 with a standard deviation of *σ*_sys,ij,target_ and added to the “true” Δ*S*_*ij*_,

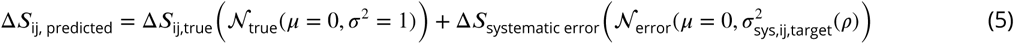

We ignore the potential complication of finite experimental error in this thought experiment, presuming the experimental uncertainty is sufficiently small as to be negligible.

The *speedup* in synthesizing molecules that reach this 10× selectivity gain threshold is calculated, as a function of *ρ*, as the ratio of the number of compounds that exceed the selectivity threshold in the case that molecules predicted to fall below this threshold by free energy calculations were triaged and not synthesized, divided by the number of compounds that exceeded the selectivity threshold without the benefit of free energy triage. This process was repeated for a 100× (2.8 kcal/mol, 2 log_10_ unit) selectivity optimization and 50 linearly spaced values of the correlation coefficient (*ρ*) between 0 and 1, for four values of *σ*_selectivity_, using a sample size of 4×10^7^ compounds.

The above model assumes that the number of compounds scored and synthesized is essentially unlimited. To assess the impact these methods might have on real drug discovery projects, where the number of compounds scored and synthesized are limited by computational and chemistry resources, we altered the above model to consider the number of compounds scored, the number of compounds triggered for synthesis, and the threshold a compound needed to reach in order to be considered for synthesis.

We repeated the mode detailed above, this time scoring only the following numbers of compounds: 10, 50, 100, 200, 500, the range from 1000 to 10000 in steps of 1000, and the range from 10000 to 100 000 in steps of 2000. Compounds were drawn from a true distribution of Δ*S*_*ij,true*_ (𝒩true(*µ* = 0, *σ*^2^ = 1)) and triaged using a free energy method as detailed above with a per-target systematic error (*σ*_sys,ij,target_) of 1 kcal/mol. The top predicted compounds that meet or surpass a synthesis rule, up to a maximum of 10 compounds, are selected for synthesis. Here, we consider synthesis rules of 100×, 500× and 1000× when trying to design 100× (2.8 kcal/mol, 2 log_10_ unit) improvements in selectivity. The *speedup* was calculated as the number of synthesized compounds whose Δ*S*_*ij,true*_ reaches the desired 100× threshold divided by the expected value (𝔼_*selective*_) for a selective compound given the number of synthesized compounds. This expectation can be calculated as,

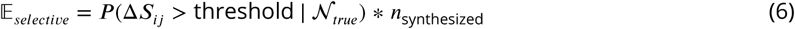

Where *P*(Δ*S*_*ij*_ *>* threshold |.𝒩_*true*_) is the probability Δ*S*_*ij,true*_ for some compound is better than a particular selectivity threshold given the distribution of Δ *S*_*ij,true*_(𝒩 _*true*_ (*µ* = 0, *σ*^2^ = 1)) for 100 000 000 compounds, and *n*_synthesized_ is the number of compounds synthesized. If no compounds were predicted to meet or surpass the synthesis rule, the speedup was assigned a default value of 1. We performed 1000 replicates of each condition and report the mean and 95 % CI in Figure1D.

### Numerical model of impact of statistical error on selectivity optimization

To model the impact of finite statistical error in the alchemical free energy calculations, a similar scheme was used with the following modifications: Each proposed compound was triaged by a free energy calculation with a per target systematic error (*σ*_sys,ij,target_) of 1.0 kcal/mol [4] and a specified correlation coefficient *ρ*. A *σ*_selectivity_ was calculated according to Equation 3, this time considering the statistical terms as non-negligible. The per target statistical error (*σ*_stat,ij,target_) was defined as,

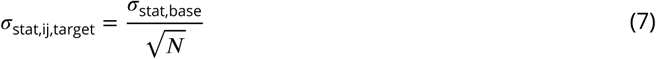

where *N* is the relative effort put into running sampling the calculation and *σ*_stat,base_ is such that when *N* = 1, *σ*_stat,ij,target_ = 0.2 kcal/mol. The statistical error is propagated assuming it is uncorrelated, as independent sets of calculations are used for each target, giving us the second set of terms in 3. This gives an updated model for the error in predicted change in selectivity Δ*S*_*ij*_. The systematic and statistical errors were modeled as Gaussian noise added to the true distribution,

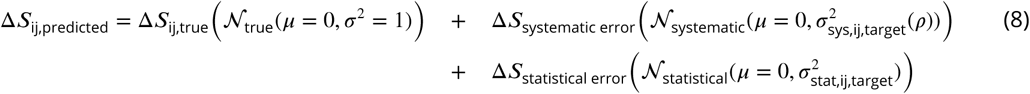

Any compound predicted to have an improvement in selectivity of above the threshold (either 1.4 kcal/mol (1 log_10_ units) or 2.8 kcal/mol (2 log_10_ units)) would then be made and have its selectivity experimentally measured, using an experimental method with perfect accuracy. The speedup value for each value of *p* is calculated as previously described.

### Binding Site Similarity analysis

To quantify the similarity between the different kinase pairs, a structure-informed binding site sequence comparison was performed. In the KLIFS database, the binding site of typical human kinases is defined by 85 residues, comprising known kinase motives (DFG, hinge, G-loop, aC-helix, …), which cover potential interactions with type I-IV inhibitors [58, 59]. KLIFS provides a multiple sequence alignment in which each kinase sequence is mapped to these 85 binding site residues. This mapping was used to calculate the sequence identify between the three kinases CDK2, CDK9, and ERK2 used in this study (Figure S1 and Table S1). The score shows the percentage of identical residues between two kinases with respect to the 85 positions.

For structural comparison, the respective pdbs of the two kinases were downloaded from the pdb (CDK2-4bci/CDK9-4bck) and CDK2-5k4j/ERK2-5k4i). PyMol v.2.3.0 was used for preprocessing and alignment of the structures. For all structures only chain A was kept. Additionally, for structure 4bck alternate location C was chosen only. Next, binding sites were selected as all residues withing 10 A of the co-crystallized ligand, yielding. Finally, the respective binding site pairs were aligned using PyMol’s default align function and the RMSD was returned. The following is an example command: `create [pdb]_bs, byres [pdb]_A within 10 of ([pdb]_A and resn [lig_name])`

### Extracting the binding free energy Δ*G* from reported experimental data

*K*_*i*_ values were derived from IC_50_ measurements reported for the ERK2/CDK2 data set (Figure 3), assuming Michaelis-Menten binding kinetics for an ATP-competitive inhibitor,

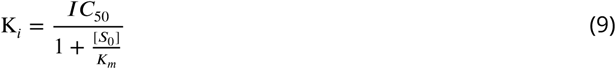

Where the Michaelis-Menten constant for ATP (*K*_*m*_ (ATP)) is much larger than the initial concentration of ATP, *S*_0_, so that IC_50_ ≈K_*i*_.

These *K*_*i*_ values were then used to calculate a Δ*G* (Equation 10),

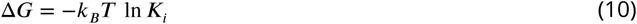

Here, *k*_*B*_ is the Boltzmann constant and *T* is absolute temperature (taken to be room temperature, *T* ∼ 300K).

For the CDK2/CDK9 data set, the authors note that the assumption *K*_*m*_ (ATP) *»* S_0_ does *not* hold, and report *K*_*i*_s derived from their *IC*_50_ measurements using the *K*_*m*_ (ATP) for each kinase, as well as the *S*_0_ from their assay. These values were then converted to Δ*G* using Equation 10. For both data sets, these derived Δ*G* were used to calculate Δ Δ*G* between ligands for each kinase target.

As mentioned above, the assumption that *K*_*m*_ (ATP) *»* S_0_ may not always hold, and changes in IC_50_ may be driven by factors other than changes in ligand binding affinity. However, these assumptions have been used successfully to estimate relative free energies previously [62, 69]. Further, data was taken from the same lab and assay for each target. By using assays with the same kinase construct and ATP concentration, the relative free energies (Δ Δ*G*_*ij*_) should be well determined for compounds within the assay. Even if the absolute free energies (Δ*G*_*i*_) are off due to uncertainties in *K*_*m*_ (ATP) or *S*_0_, they will be off by the same constant, which will cancel when calculating Δ Δ*G*_*ij*_.

### Structure Preparation

Structures from the Shao [53] (CDK2/CDK9), Hole [60] (CDK2/CDK9), and Blake [54] (CDK2/ERK2) papers were downloaded from the PDB [70], selecting structures with the same co-ligand crystallized.

For the Shao (CDK2/CDK9) data set, PDB IDs 4BCK (CDK2) and 4BCI (CDK9) were selected, which have ligand 12c cocrystallized. For the Blake data set (ERK2/CDK2), 5K4J (CDK2) and 5K4I (ERK2) were selected, cocrystallized with ligand 21. The structures were prepared using Schrodinger’s Protein Preparation Wizard [71] (Maestro, Release 2017-3). This pipeline modeled in internal loops and missing atoms, added hydrogens at the reported experimental pH (7.0 for the Shao data set, 7.3 for the Blake data set) for both the protein and the ligand. All crystal waters were retained. The ligand was assigned protonation and tautomer states using Epik at the experimental pH±2, and hydrogen bonding was optimized using PROPKA at the experimental pH±2. Finally, the entire structure was minimized using OPLS3 with an RMSD cutoff of 0.3Å.

### Ligand Pose Generation

Ligands were extracted from the publication entries in the BindingDB as 2D SMILES strings. 3D conformations were generated using LigPrep with OPLS3 [4]. Ionization state was assigned using Epik at experimental pH±2. Stereoisomers were computed by retaining any specified chiralities and varying the rest. The tautomer and ionization state with the lowest Epik state penalty was selected for use in the calculation. Any ligands predicted to have a positive or negative charge in its lowest Epik state penalty was excluded, with the exception of Compound 9 from the Blake data set. This ligand was predicted to have a +1 charge for its lowest state penalty state. The neutral form the ligand was include for the sake of cycle closure in the FEP+ map, but was ignored for the sake any analysis afterwards. Ligand poses were generated by first aligning to the co-crystal ligand using the Largest Common Bemis-Murcko scaffold with fuzzy matching (Maestro, Release 2017-3). Ligands that were poorly aligned or failed to align were then aligned using Maximum Common Substructure (MCSS). Finally, large R-groups conformations were sampled with MM-GBSA using a common core restraint, VSGB solvation model, and OPLS3 force field. No flexible residues were defined for the protein.

### Free Energy Calculations

The FEP+ panel (Maestro, Release 2017-3) was used to generate perturbation maps. FEP+ calculations were run using the FEP+ panel from Maestro release 2018-3 in order to take advantage of the newest force field (OPLS3e) parameters available at the time. Any missing ligand torsions were fit using the automated FFbuilder protocol [7]. Custom charges were assigned using the OPLS3e force field using input geometries, according to the automated FEP+ workflow in Maestro Release 2018-3. Neutral perturbations were run for 15 ns per replica, using an NPT ensemble and water buffer size of 5Å. The SPC water model was used. A GCMC solvation protocol was used to sample buried water molecules in the binding pocket prior to the calculation, which discards any retained crystal waters.

### Statistical Analysis of FEP+ calculations

To quantify the overall errors in the FEP+ calculations, we computed the mean unsigned error (MUE),

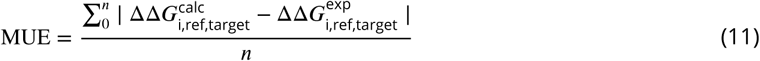

and the root mean squared error (RMSE)

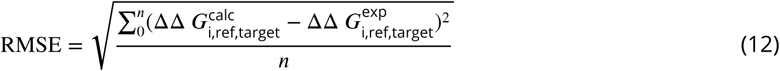

MUE and RMSE were computed for Δ Δ*G*_*ij*,target_. For each ligand *i*, Δ Δ*G*_*i,ref*,target_ is defined where ref is a reference compound.

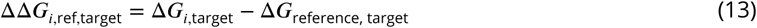

For the CDK2/CDK9 data set, compound 1a was used as the reference compound, as it was the first compound from which the others in the series were derived. For the CDK2/ERK2 data set, compound 6 was used as the reference compound, since it was the compound from which the investigation was launch. A metabolite of compound 6 (not included in the data set here) was used as the starting compound from which the rest were derived. To account for the finite ligand sample size, we used 10 000 replicates of bootstrapping with replacement to estimate 95% confidence intervals. The code used to bootstrap these values is available on GitHub [https://github.com/choderalab/selectivity].

To compute the per-target statistical error (*σ*_stat,ij,target_) for each *i*,ref pair of ligands, we used the standard deviation of 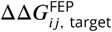, where *j* is the reference compound, from the Bayesian model described in depth below in the **Methods** section. To compute the per target systematic error (*σ*_sys,ij,target_), we calculated the mean of *ϵ*_*ij*,target_, where *j* is the reference compound, described in equation 21 in the Bayesian Model section of the **Methods**.

### Quantification of the correlation coefficient *σ*

To quantify *ρ*, we built a Bayesian graphical model using pymc3 3.5 [72] and theano 1.0.3 [73]. All code for this model is available on GitHub [https://github.com/choderalab/selectivity].

For each phase (complex and solvent), the prior for the absolute free energy (*G*) of ligand *i* (up to an arbitrary additive constant for each thermodynamic phase, ligand-in-complex or ligand-in-solvent), was treated as a normal distribution (Equation 15).

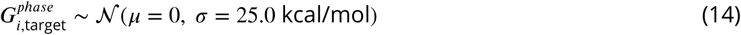

To improve sampling efficiency, for each phase, one ligand was chosen as the reference, and pinned to an absolute free energy of *G* = 0, with a standard deviation of 1 kcal/mol.

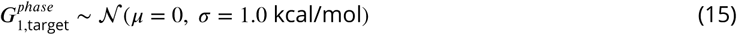

For each edge of the FEP map (ligand *i* –> ligand *j*), there is a contribution from dummy atoms, that was modeled as in Equation 16. Note that here, unlike what was done in Figure 4, ligand *j* is not necessarily a reference compound.

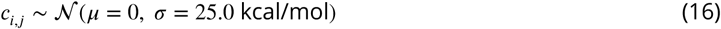

The model was conditioned by including data from the FEP+ calculation.

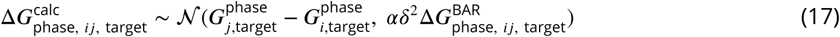

where 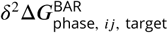 is the reported BAR uncertainty from the calculation, and 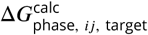 is the BAR estimate of the free energy for the perturbation between ligands *i* and *j* in a given phase. *α* is a scaling parameter shared by all 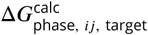 for each target. Such scaling is necessary to account for the BAR statistical uncertainty underestimating cycle closure statistical uncertainty of our calculations, shown by Figure S8.

From this, we can calculate the 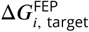 for each ligand and target,

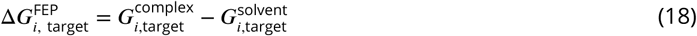

From 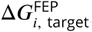, we calculated 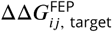 for each pair of ligands, filtering out pairs where *i* and *j* are the same ligand and where the reciprocal was already included.

The experimental binding affinity was treated as a true value 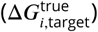 corrupted by experimental uncertainty, which is assumed to be 0.3 kcal/mol [6]. There are a number of studies that report on the reproducibility and uncertainty of intra-lab IC_50_ measurements, ranging from as small as 0.22 kcal/mol [62] to as high as 0.4 kcal/mol [6]. The assumed value falls within this range and is in good agreement with the uncertainty reported from multiple replicate measurements in internal data sets at Novartis [63].

The values reported in the papers 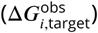 were treated as observations from this distribution (Equation 19),

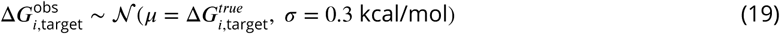

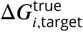 was assigned a weak normal prior, as in Equation 20,

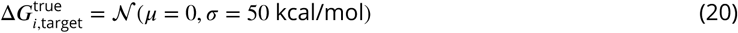

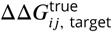 for each pair of ligands was calculated from 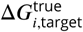, filtering out pairs where *i* and *j* are the same ligand and where the reciprocal was already included as above.

The error for a given ligand was calculated as

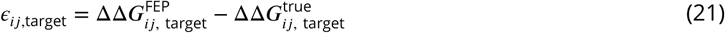

From these *ϵ* values, we calculated the correlation coefficient, *ρ*, from the sampled errors for the finite set of molecules for which measurements were available,

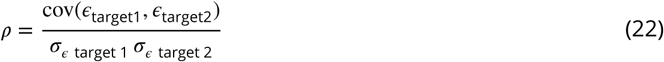

where *σ* _*ϵ* target 2_ is the standard deviation of *ϵ*_ij,target_.

To quantify *ρ* from these calculations, the default NUTS sampler with `jitter+adapt_diag` initialization, 3 000 tuning steps, and the default target accept probability was used to draw 20 000 samples from the model.

### Calculating the marginal distribution of speedup

To quantify the expected speedup from the calculations we ran, we utilized 10^4^ replicates of the scheme detailed above to calculate the speedup given parameters *ρ, σ*_sys,ij,1_, and *σ*_sys,ij,2_, in the regime of infinite effort and zero statistical error. Using Numpy 1.14.2, *ρ* was drawn from a normal distribution with the mean and standard deviation from the posterior distribution of *ρ* from the Bayesian Graphical model. The per-target systematic errors, *σ*_sys,ij,1_ and *σ*_sys,ij,2_, were estimated from the mean of the absolute value of *ϵ*_ij,1_ and *ϵ*_ij,2_, which are the magnitude of errors from the Bayesian graphical model. *σ*_selectivity_ was calculated using Equation 3. 10^6^ molecules were proposed from true normal distribution, as above. The error of the computational method was modeled as in Equation 5.

## Data Availability

All curated starting structures, FEP+ results, and data analysis scripts and notebooks are available on GitHub: https://github.com/choderalab/selectivity

## Acknowledgments

The authors are grateful to Patrick Grinaway (ORCID: 0000-0002-9762-4201) for useful discussions about Bayesian statistics and Mehtap Işık (ORCID: 0000-0002-6789-952X) for useful discussion about kinase inhibitor protonation states. SKA is grateful to Haoyu S. Yu, Wei Chen, and Dmitry Lupyan for advice on running FEP+ calculations.

## Author Contributions

Conceptualization: SKA, LW, RA, JDC; Methodology: SKA, LW, JDC; Formal Analysis: SKA, JDC, LW; Data Curation: SKA, SP; Investigation: SKA, SP, AV; Writing – Original Draft: SKA, JDC; Writing – Review & Editing: SKA, JDC, LW, AV, RA; Visualization: SKA, JDC, LW; Supervision: LW, JDC, RA; Project Administration: SKA, LW, JDC, RA; Funding Acquisition: RA, JDC; Resources: LW, JDC

