## Supplemental information for "Is structure based drug design ready for selectivity optimization?"

**Table S1. The CDK2 and CDK9 binding sites are more similar than the CDK2 and ERK2 binding sites**
Sequence based similarity of the binding sites based on multiple sequence alignments of the 85 residues annotated by the KLIFS. Upper triangle shows the ratio of sequence identity, lower triangle, the number of matching residues out of 85 binding site residues. database [1, 2]

| Kinase | CDK2 | CDK9 | ERK2 |
|---|---|---|---|
| CDK2 | 1.0 | 0.57 | 0.52 |
| CDK9 | 47 | 1.0 | 0.52 |
| ERK2 | 43 | 43 | 1.0 |

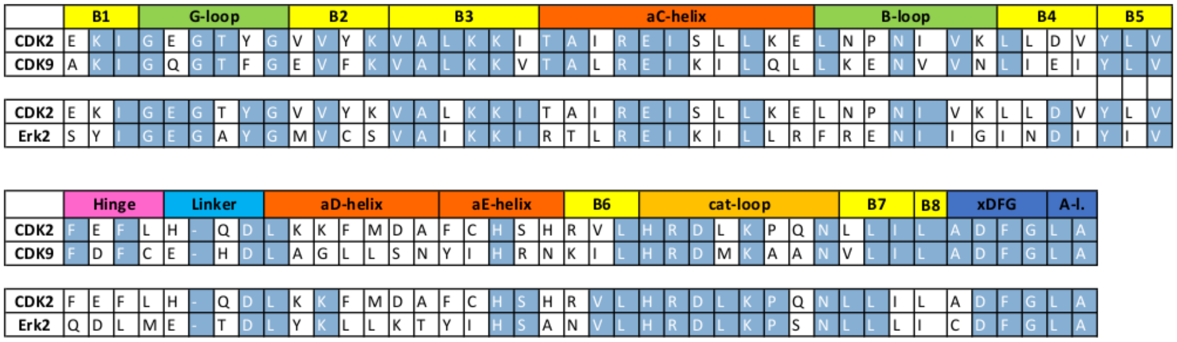


**Figure S1. Sequence identity of the kinase pairs by binding site motif**
Binding site sequence identity for CDK2/CDK9 and CDK2/ERK2 by binding site motif, as defined by the KLIFs database [1, 2]. Identical residues between the pairs are shown in blue.

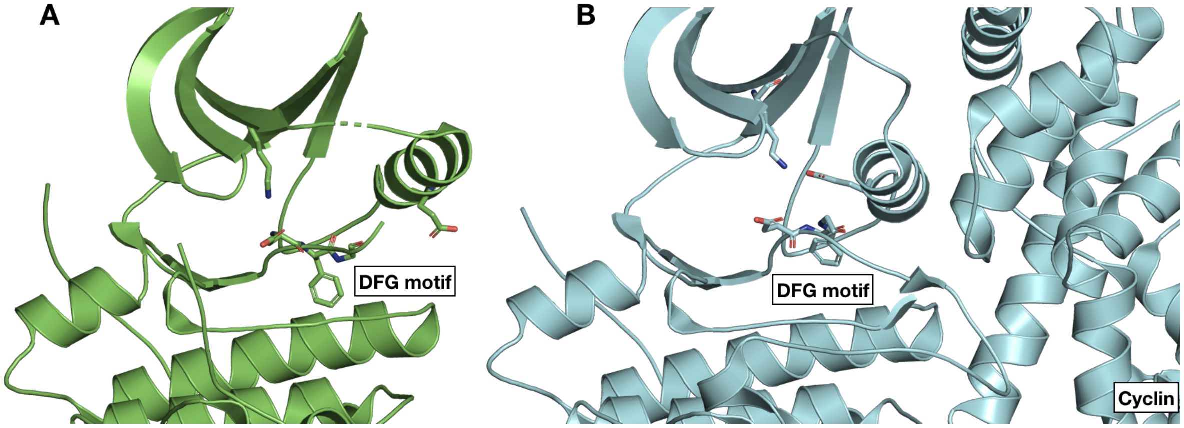


**Figure S2. CDK2 adopts an inactive conformation in the crystal structure used for the CDK2/ERK2 calculations**
(**A**) CDK2 (5K4J) adopts an inactive conformation in the absence of its cyclin. The DFG motif is in a DFG-in conformation, with the $\alpha$C helix rotated outwards, breaking the salt bridge between K33 and E51 (Uniprot numbering) that is typically a marker of an active conformation. Notably, the Phe in the DFG motif does not completely form the hydrophobic spine due to the rotation of the $\alpha$C helix [3] (**B**) The CDK2 structure used for the CDK2/CDK9 calculations (4BCK) contains cyclin A and adopts a DFG-in/$\alpha$C helix-in conformation that forms the salt bridge between K33 and E51. This is typically indicative of a fully active kinase [4, 5].

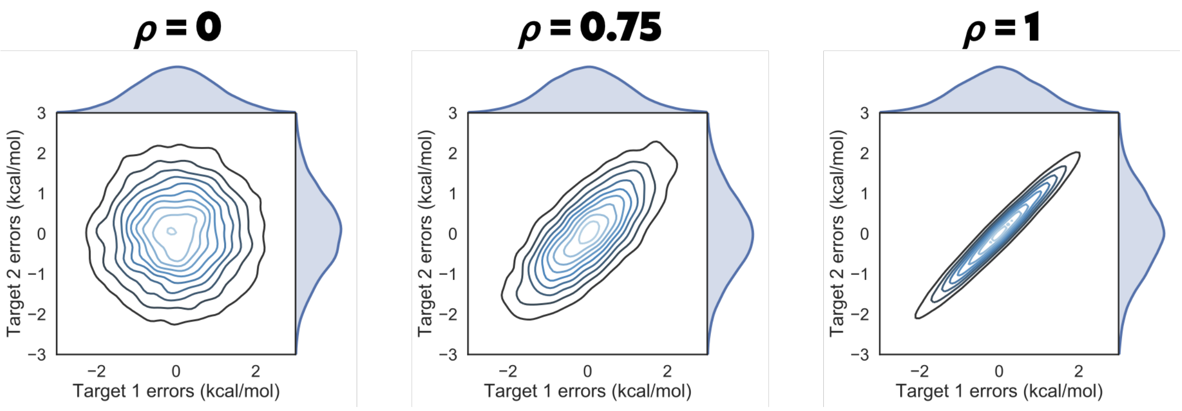


**Figure S3. Correlation coefficient $\rho$ controls the shape of the joint marginal distribution of errors**
As $\rho$ increases, the joint marginal distribution of errors become more diagonal. Each panel shows 10000 samples drawn from a multivariate normal distribution centered around 0 kcal/mol, where the per target error was set to 1 kcal/mol and $\rho$ to the value indicated in bold over the plot.

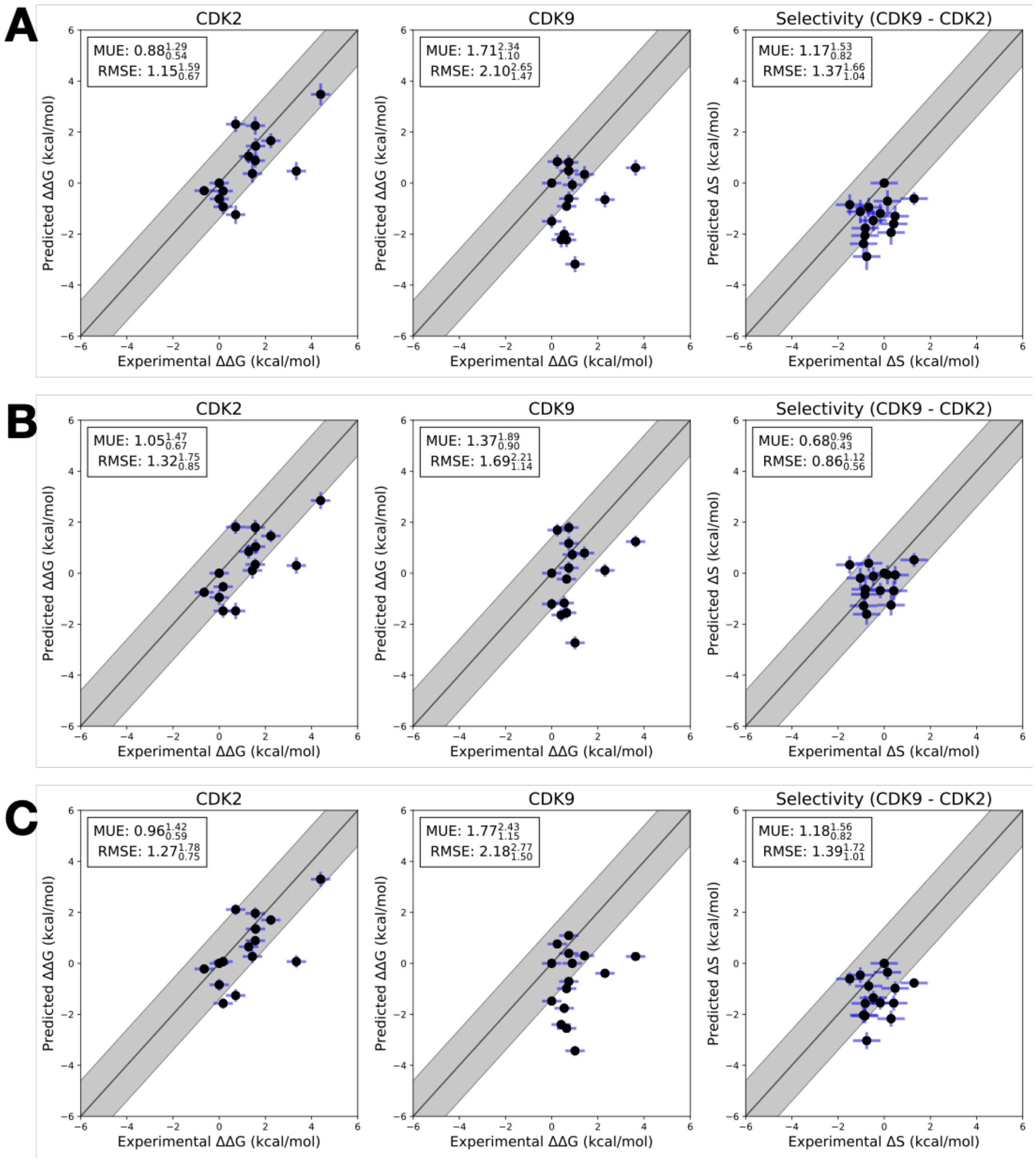


**Figure S4. Each replicate of the CDK2/CDK9 calculations yields a consistent RMSE and MUE**
Three replicates of the CDK2/CDK9 calculations with different random seeds, but otherwise the same input structures, files, and parameters. The experimental values are shown on the X-axis and calculated values on the Y-axis. Each data point corresponds to a transformation between a ligand $i$ to a set reference ligand $j$ (Compound 1a) for a given target. All values are shown in units of kcal/mol. The blue vertical error bars are $\sigma_{\mathrm{stat,ij,target}}$, which was estimated by calculating the standard deviation of $\Delta\Delta G^{\mathrm{FEP}}_{ij,\,\mathrm{target}}$ from the Bayesian model described in depth in **Methods**. The horizontal error bars show $\delta\Delta\Delta G^{exp}_{ij}$ based on the assumed uncertainty of 0.3 kcal/mol[6, 7] for each $\Delta G^{exp}_{i}$ expanded assuming no correlation between each measurement. For selectivity, the errors were propagated under the assumption that they were completely uncorrelated. The black line indicates agreement between calculation and experiment, while the gray shaded region represent 1.36 kcal/mol (or 1 log unit) error. The MUE and RMSE are shown on each plot with bootstrapped 95% confidence intervals. (**A**) Replicate 1 (**B**) Replicate 2 (**C**) Replicate 3

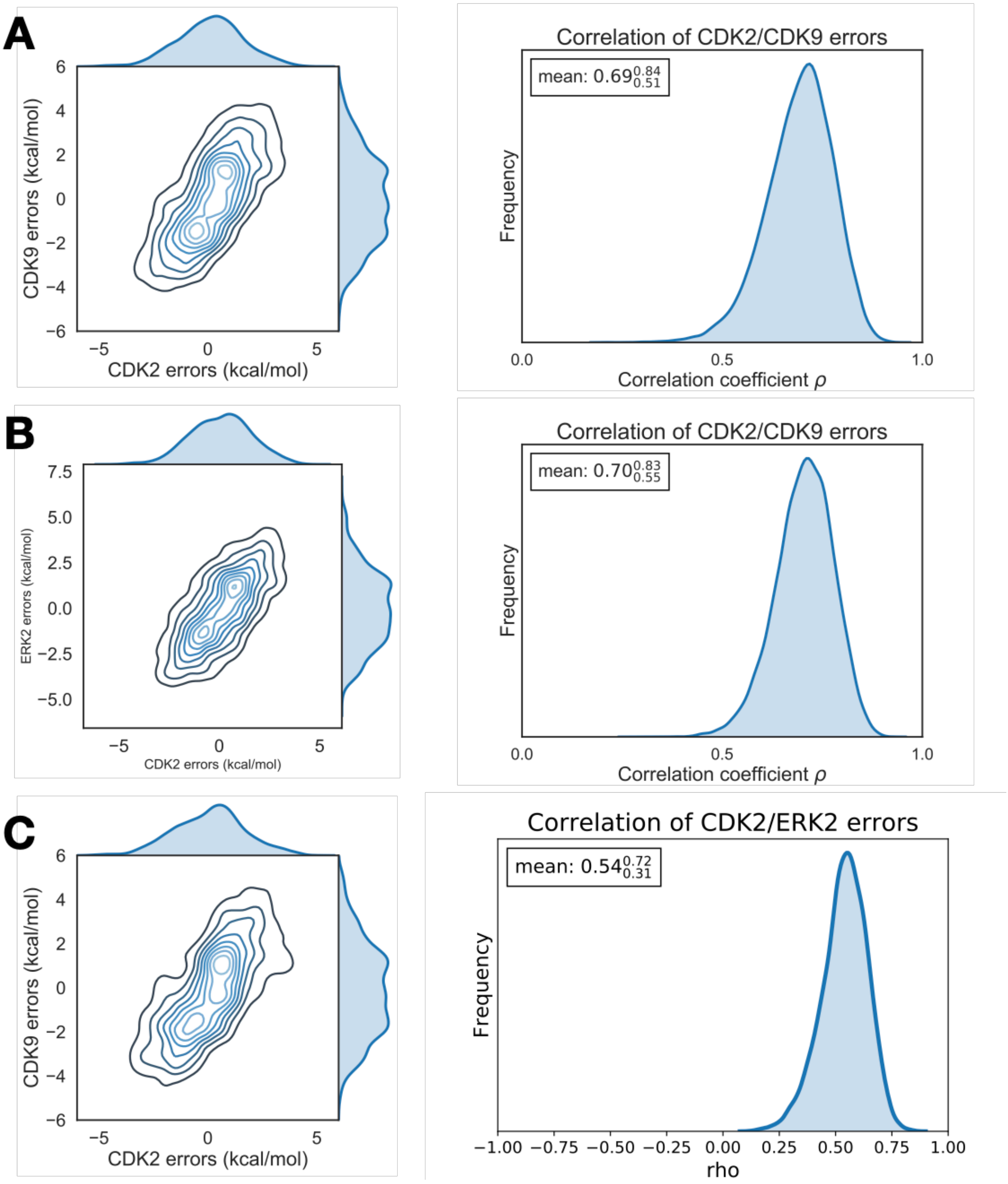


**Figure S5. Each replicate of the CDK2/CDK9 calculations yields consistent errors and correlation coefficient**
(**A**) (*left*) The joint posterior distribution of the prediction errors for CDK2 (X-axis) and CDK9 (Y-axis) from the Bayesian graphical model for replicate 1. (*right*) The posterior marginal distribution of the correlation coefficient ($\rho$) is shown in gray for replicate 1. The inserted box shows the mean and 95% confidence interval for the correlation coefficient. (**B**) and (**C**) The same as above, but for replicates 2 and 3, respectively

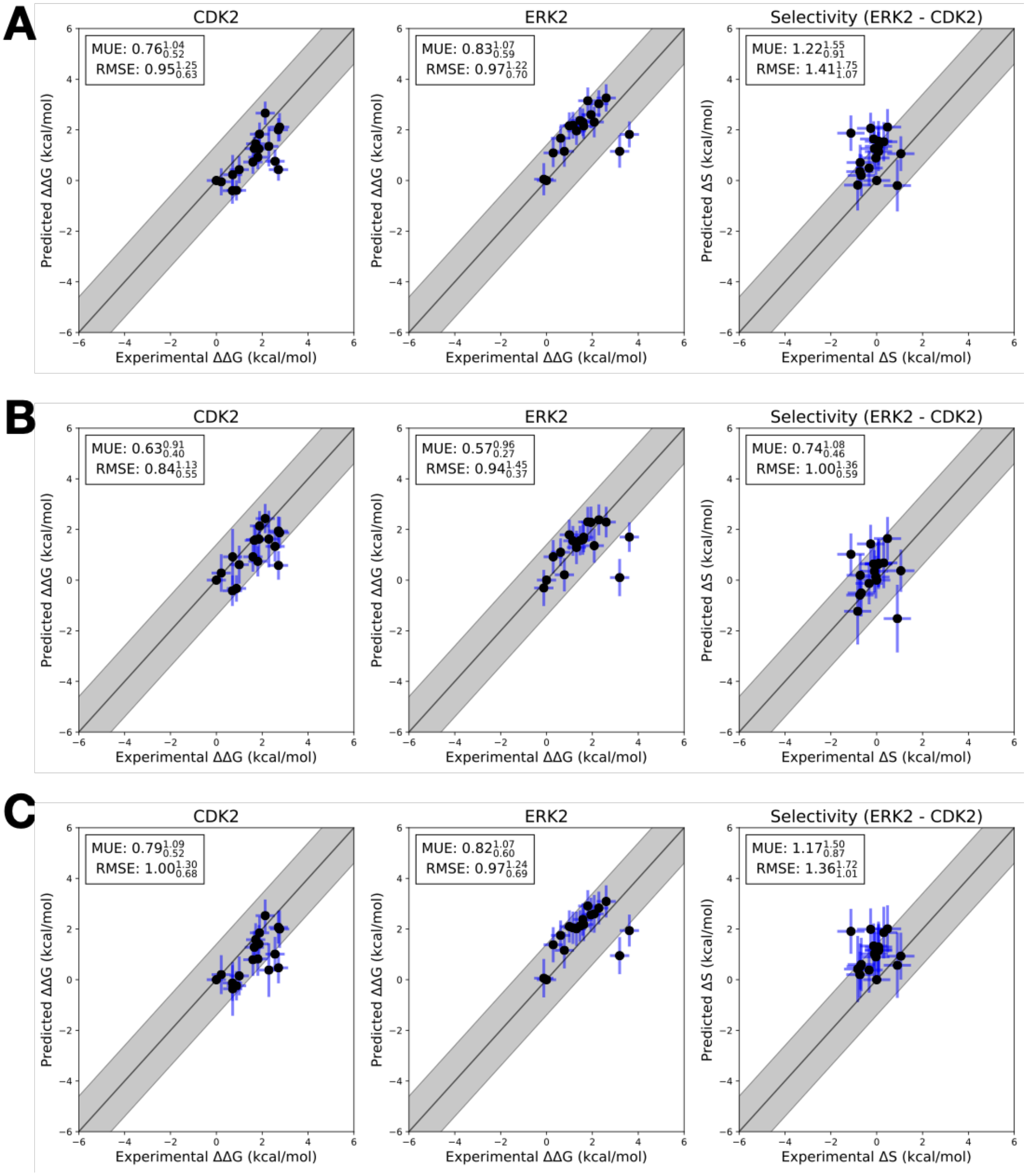


**Figure S6. Each replicate of the CDK2/ERK2 calculations yields a consistent RMSE and MUE**
Three replicates of the CDK2/ERK2 calculations with different random seeds, but otherwise the same input structures, files, and parameters. The experimental values are shown on the X-axis and calculated values on the Y-axis. Each data point corresponds to a transformation between a ligand $i$ to reference ligand $j$ (Compound 6) for a given target. All values are shown in units of kcal/mol. The blue vertical error bars are $\sigma_{\text{stat,ij,target}}$, which was estimated by calculating the standard deviation of $\Delta\Delta G^{\text{FEP}}_{ij,\text{ target}}$ from the Bayesian model described in depth in **Methods**. The horizontal error bars show $\delta\Delta\Delta G^{exp}_{ij}$ based on the assumed uncertainty of 0.3 kcal/mol[6, 7] for each $\Delta G^{exp}_{i}$ expanded assuming no correlation between each measurement. For selectivity, the errors were propagated under the assumption that they were completely uncorrelated. The black line indicates agreement between calculation and experiment, while the gray shaded region represent 1.36 kcal/mol (or 1 log unit) error. The MUE and RMSE are shown on each plot with bootstrapped 95% confidence intervals. (**A**) Replicate 1 (**B**) Replicate 2 (**C**) Replicate 3

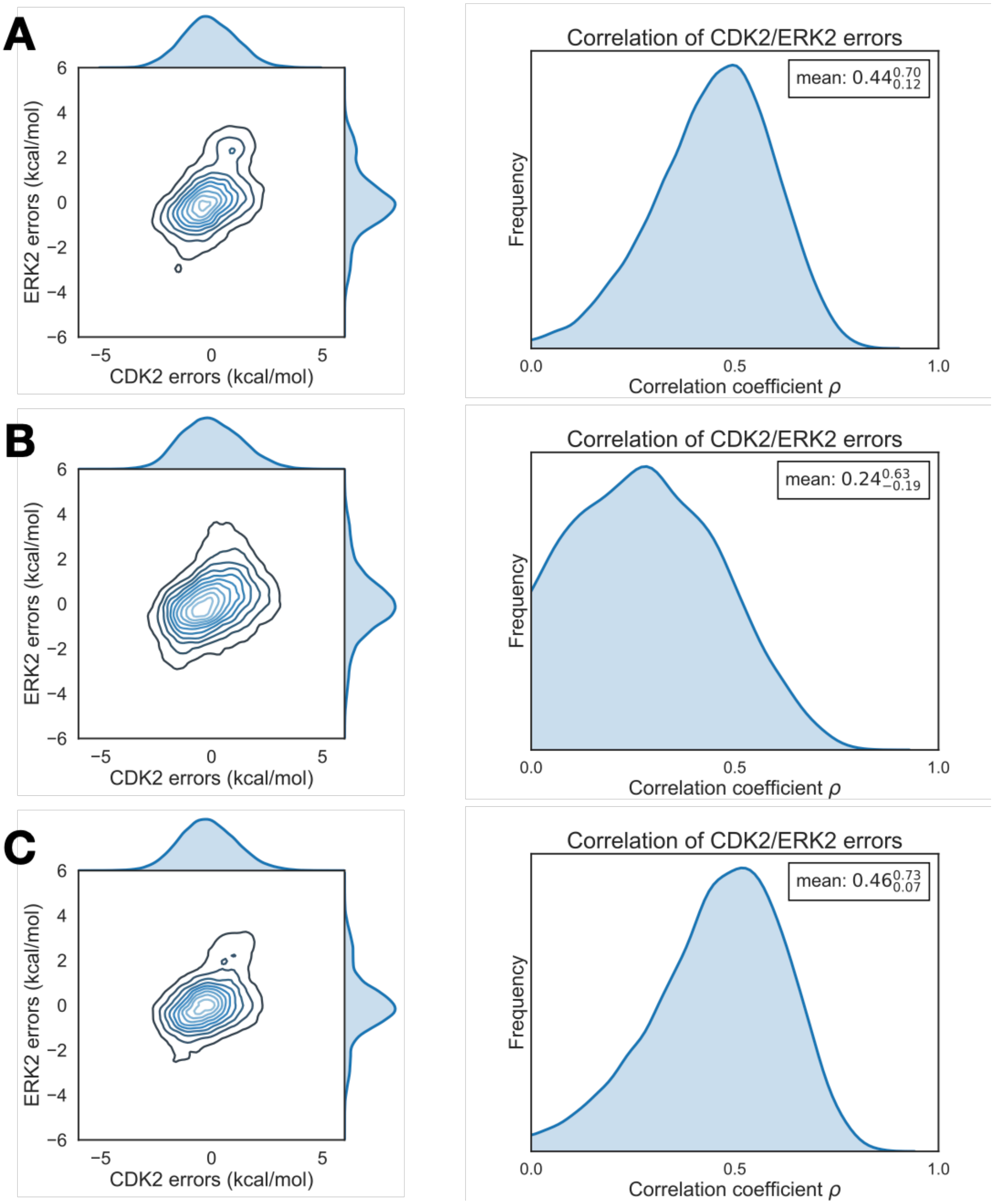


**Figure S7. Each replicate of the CDK2/ERK2 calculations yields consistent errors and correlation coefficient** (**A**) (*left*) The joint posterior distribution of the prediction errors for CDK2 (X-axis) and ERK2 (Y-axis) from the Bayesian graphical model for replicate 1. (*right*) The posterior marginal distribution of the correlation coefficient ($\rho$) is shown in gray for replcicate 1. The inserted box shows the mean and 95% confidence interval for the correlation coefficient. (**B**) and (**C**) The same as above, but for replicates 2 and 3, respectively

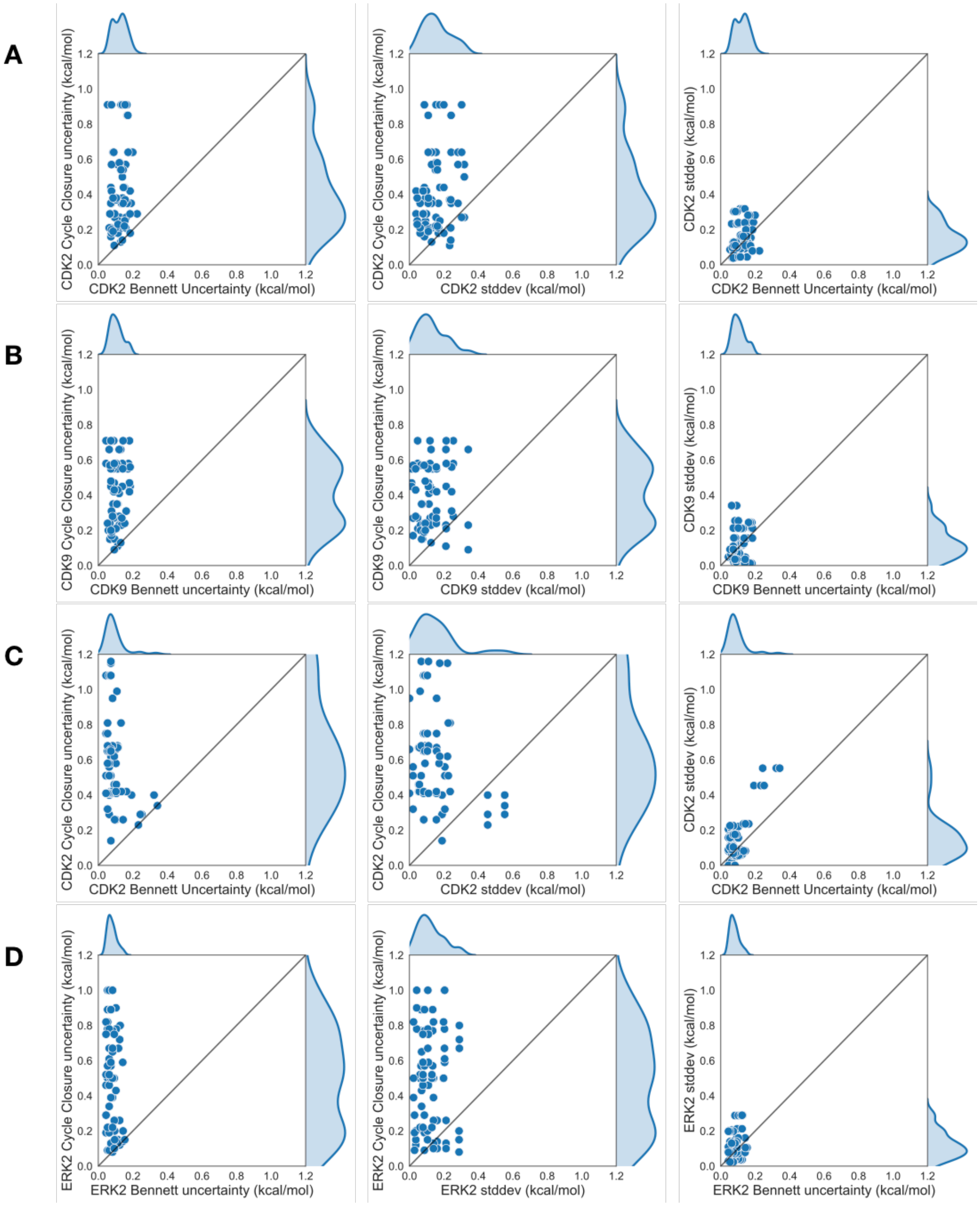


**Figure S8. The standard deviation and Bennett error for each edge is smaller than the estimated cycle closure uncertainties**
Pairwise Comparisons of the cycle closure uncertainty, the Bennett uncertainty, and the standard deviation of three replicate calculations, reported in kcal/mol. Each point corresponds to an edge of the FEP map. The edges for all three replicates are pooled and shown together. (**A and B**) CDK2/CDK9 calculations from the Shao data set (**C and D**) CDK2/ERK2 calculations from the Blake data set

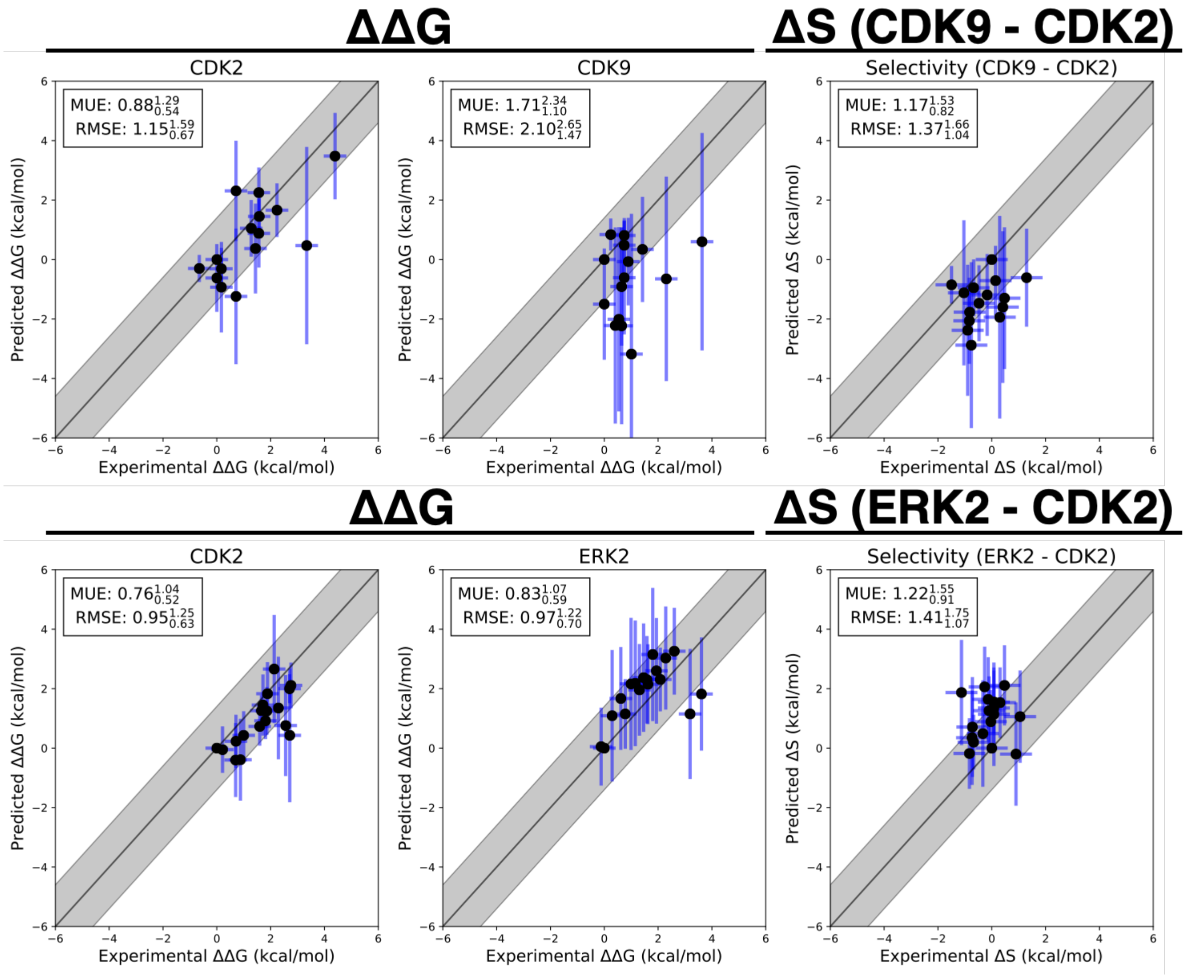


**Figure S9. The statistical and systematic error explain deviation from experimental measurements**
$\Delta\Delta G_{ij,\mathrm{target}}$ and $\Delta S_{ij}$ predictions for CDK2/ERK2 from the Blake data sets (*top*), and CDK2/CDK9 (*bottom*) from the Shao data sets. The experimental values are shown on the X-axis and calculated values on the Y-axis. Each data point corresponds to a transformation between a ligand $i$ to a set reference ligand $j$ for a given target. All values are shown in units of kcal/mol. The horizontal error bars show the $\delta\Delta\Delta G_{ij}^{exp}$ based on the assumed uncertainty of 0.3 kcal/mol[6, 7] for each $\Delta G_i^{exp}$. We show the estimated error ($\sigma_{\mathrm{stat,ij,target}}$ + $\sigma_{\mathrm{sys,ij,target}}$) as vertical blue error bars, which are one standard error. $\sigma_{\mathrm{stat,ij,target}}$ was estimated by calculating the standard deviation of $\Delta\Delta G^{\mathrm{FEP}}_{ij,\,\mathrm{target}}$ from the Bayesian model described in depth in **Methods**. $\sigma_{\mathrm{sys,ij,target}}$ was estimated from the mean of $\epsilon_{ij,\mathrm{target}}$ described in equation 15. For the $\Delta S$ panels, $\sigma_{\mathrm{selectivity}}$ (vertical blue error bars) was calculated according to Equation **??** using the estimated correlation coefficients from Figure 6.The black line indicates agreement between calculation and experiment, while the gray shaded region represent 1.36 kcal/mol (or 1 log unit) error. The MUE and RMSE are shown on each plot with bootstrapped 95% confidence intervals.

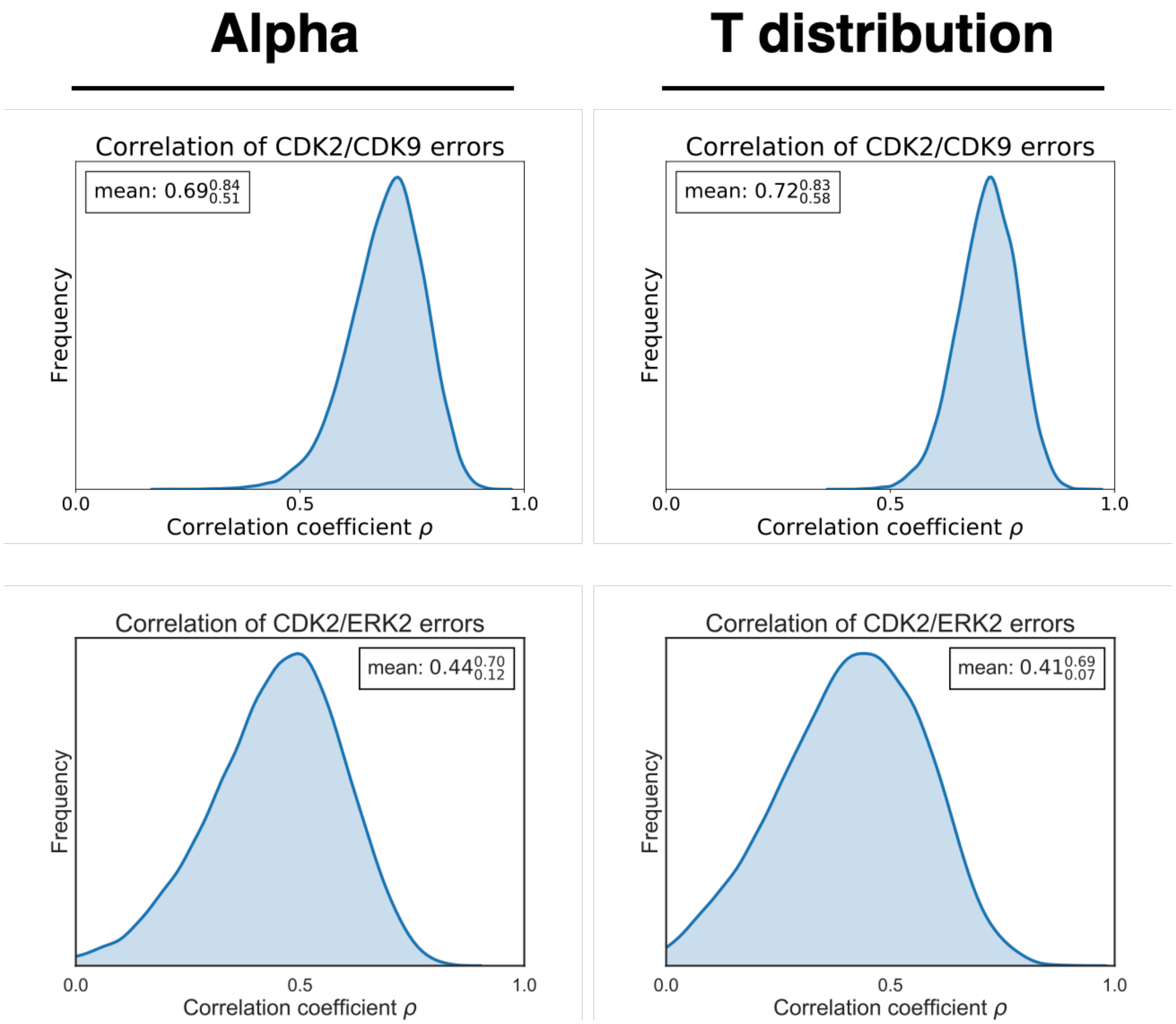


**Figure S10. Estimates of correlation coefficient $\rho$ are insensitive to use of scaling term $\alpha$ or Student's t-distribution** (*left*) Estimate of correlation coefficient $\rho$ for replicate 1 of the CDK2/CDK9 (*top*) and CDK2/ERK2 (*bottom*) calculations using a scaling term ($\alpha$) to account for the BAR error underestimating the cycle closure statistical error, shown in greater detail in the **Methods** section Equation 10. (*right*) Estimate of correlation coefficient $\rho$ for CDK2/CDK9 (*top*) and CDK2/ERK2 (*bottom*) using a Student's t-distribution instead of a Normal distribution and scaling term in Equation 10
